# Assessment of community efforts to advance computational prediction of protein-protein interactions

**DOI:** 10.1101/2021.09.22.461292

**Authors:** Xu-Wen Wang, Lorenzo Madeddu, Kerstin Spirohn, Leonardo Martini, Adriano Fazzone, Luca Becchetti, Thomas P. Wytock, István A. Kovács, Olivér M. Balogh, Bettina Benczik, Mátyás Pétervári, Bence Ágg, Péter Ferdinandy, Loan Vulliard, Jörg Menche, Stefania Colonnese, Manuela Petti, Gaetano Scarano, Francesca Cuomo, Tong Hao, Florent Laval, Luc Willems, Jean-Claude Twizere, Michael A. Calderwood, Enrico Petrillo, Albert-László Barabási, Edwin K. Silverman, Joseph Loscalzo, Paola Velardi, Yang-Yu Liu

## Abstract

Comprehensive insights from the human protein-protein interaction (PPI) network, known as the human interactome, can provide important insights into the molecular mechanisms of complex biological processes and diseases. Despite the remarkable experimental efforts undertaken to date to determine the structure of the human interactome, many PPIs remain unmapped. Computational approaches, especially network-based methods, can facilitate the identification of new PPIs. Many such approaches have been proposed. However, a systematic evaluation of existing network-based methods in predicting PPIs is still lacking. Here, we report community efforts initiated by the International Network Medicine Consortium to benchmark the ability of 24 representative network-based methods to predict PPIs across five different interactomes, including a synthetic interactome generated by the duplication-mutation-complementation model, and the interactomes of four different organisms: *A. thaliana*, *C. elegans*, *S. cerevisiae*, and *H. sapiens*. We selected the top-seven methods through a computational validation on the human interactome. We next experimentally validated their top-500 predicted PPIs (in total 3,276 predicted PPIs) using the yeast two-hybrid assay, finding 1,177 new human PPIs (involving 633 proteins). Our results indicate that task-tailored similarity-based methods, which leverage the underlying network characteristics of PPIs, show superior performance over other general link prediction methods. Through experimental validation, we confirmed that the top-ranking methods show promising performance externally. For example, from the top 500 PPIs predicted by an advanced similarity-base method [MPS(B&T)], 430 were successfully tested by Y2H with 376 testing positive, yielding a precision of 87.4%. These results establish advanced similarity-based methods as powerful tools for the prediction of human PPIs.

## INTRODUCTION

Comprehensive understanding of the human PPI network (also known as the interactome) could offer global insights into cellular organization, genome function, and genotype–phenotype relationships^1,2^. Discovery of new PPIs could facilitate important interventional goals, as well, such as drug target identification and therapeutic design^3^. Despite remarkable experimental efforts in high-throughput mapping, the human interactome map remains sparse and incomplete^2,4^, and is subject to noise and investigative biases^2^. These factors represent a severe limitation to accurately understanding cellular organization and genome function. Computational methods can accelerate knowledge acquisition in biomedical networks by significantly reducing the number of alternatives to be confirmed in bench experiments^5–9^. Yet, high incompleteness of the human interactome map may reduce the effectiveness of state-of-the-art computational methods. In this context, the computational prediction of new PPIs based on experimentally observed PPIs becomes a particularly challenging but potentially highly rewarding task.

Recognizing the advantages and limitations of different computational methods in the context of PPI prediction is critical, providing the insight required to select the best predictive strategy^10–14^. To accelerate this process, the International Network Medicine Consortium initiated a project to systematically benchmark 24 representative network-based computational methods in PPI prediction through standardized performance metrics and standardized, unbiased interactome analysis (see Fig.1 for a summary of the workflow of this project). The 24 methods, as listed in Table 1, cover major categories of link prediction from similarity-based methods to probabilistic methods, factorization-based methods, diffusion-based methods, and machine learning-based methods (Fig.2). The benchmark interactomes used include those of four different organisms, *A. thaliana*^15^, *C. elegans*^16^, *S. cerevisiae*^17^, *H. sapiens*^4^, and a synthetic interactome generated by the duplication-mutation-complementation model^18^. For each of the five interactomes, we first performed 10-fold cross-validation to evaluate four performance metrics of the 24 different methods (here we refer to this process as “computational validation”). This analysis allowed us to rank the methods. Next, the top-seven methods were selected based on their performance in predicting interactions in the human interactome, and their top-500 predicted human PPIs (yielding a cumulative of 3,276 PPIs) were chosen for a systematic and unbiased experimental validation through yeast two-hybrid (Y2H) assays. In total, we validated 1,177 new PPIs involving 633 human proteins. To the best of our knowledge, no other consortium-based evaluations of PPI prediction algorithms in the literature incorporated such a large experimental validation effort.

**Figure 1:**
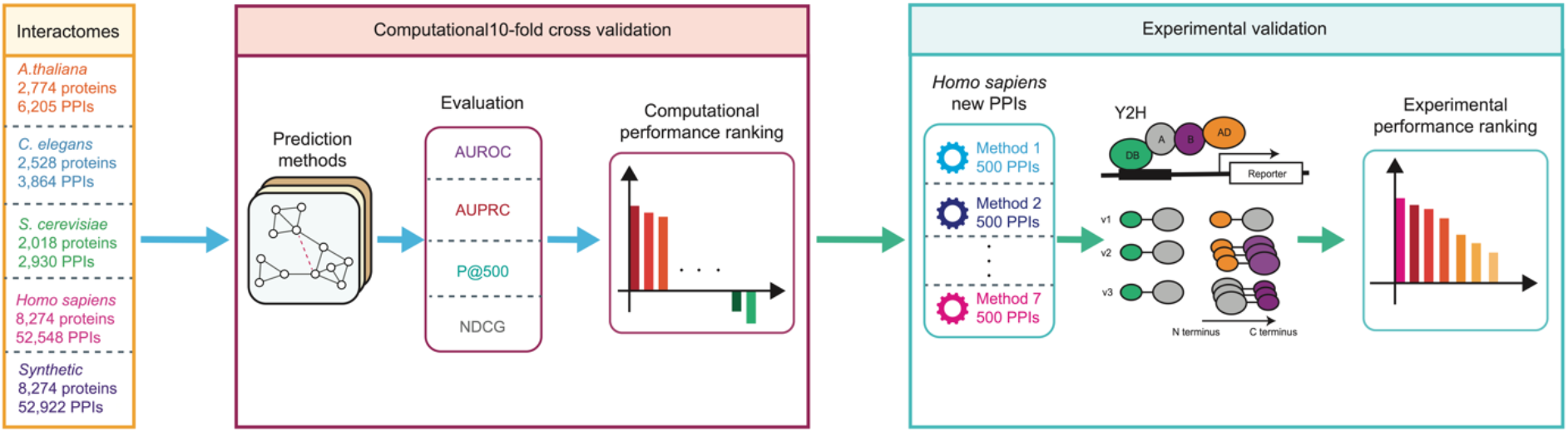
Workflow of the INMC PPI prediction project. Methods were evaluated as follows: (1) Participants were required to predict the PPIs of an *in silico* interactome (Synthetic PPI) and interactome from four different organisms: *A. thaliana*^15^, *C. elegans*^16^, *S. cerevisiae*^17^ and *H. sapiens*^4^. (2) The PPIs of each interactome were divided into training set and validation set through 10-fold cross validation. (3) Each method was evaluated using four standard metrics: Area Under the Receiver Operating Characteristic (ROC) curve (AUROC), Area Under the Precision-Recall Curve (AUPRC), Precision of the top-500 predicted PPIs (P@500), and Normalized Discounted Cumulative Gain (NDCG). (4) For the human interactome, an overall score was defined as the sum of z-scores of three metrics (AUPRC, P@500 and NDCG), after which the final rank was obtained by sorting these scores. (5) For each of the top-seven ranking methods, the top500 predicted PPIs of each method were validated experimentally. PPI: protein-protein interactions; Y2H: yeast two hybrid assay; v1-v3: assay 1-assay 3.

**Table 1.**
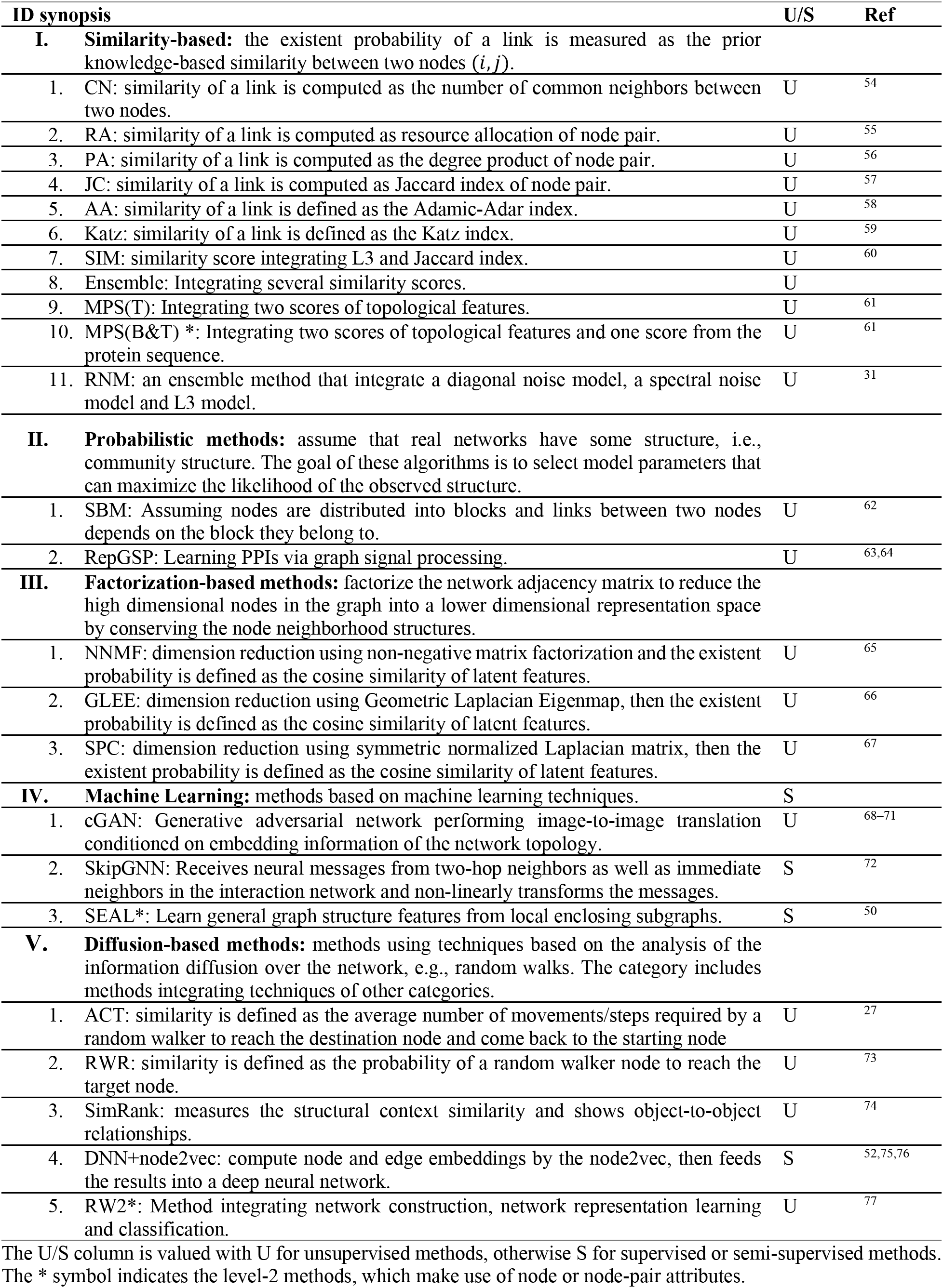
Computational Methods tested in the INMC PPI prediction project.

**Figure 2:**
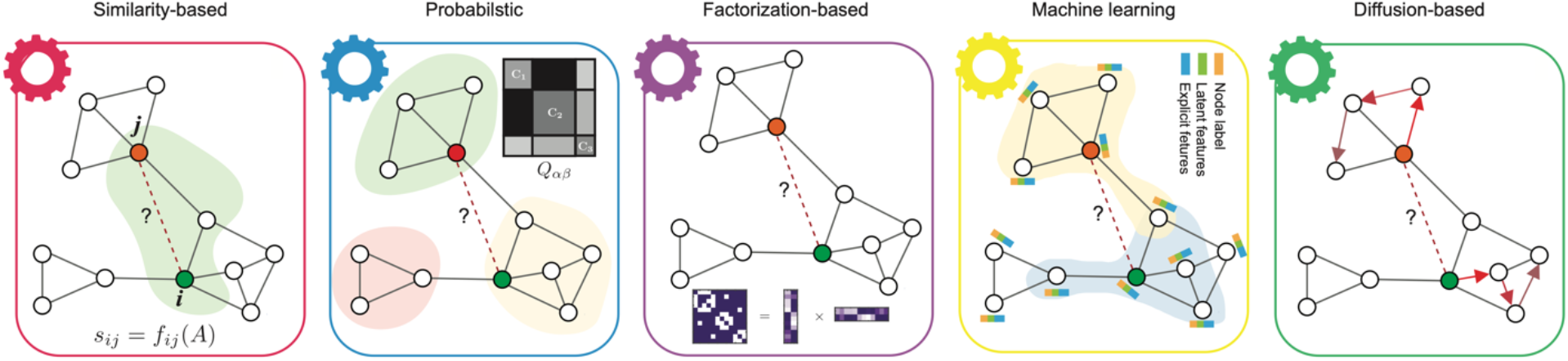
Diagram of the five major categories of link prediction methods. (1) Similarity-based methods. These methods quantify the likelihood of links based on predefined similarity functions among nodes in the graph, i.e., the common neighbors (green area). (2) Probabilistic methods. These methods assume that real networks have some structure, e.g., community structure. The goal of these algorithms is to select model parameters that can maximize the likelihood of the observed structure. The connecting probability of nodes within a community is higher than that between different communities (gray matrix). (3) Factorization-based: The goal of these methods is to learn a lower dimensional representation for each node in the graph by preserving the global network patterns. Next, the compressed representation is leveraged to predict unobserved PPIs by either calculating a similarity function or training a classifier. (4) Machine learning: There are numerous methods among machine learning categories; here, we illustrate this category using the state-of-the-art graph neural networks (GNN). Those methods embed node information by aggregating the node features, link features and graph structure using a neural network and passing the information through links in the graph. Thereafter, the learned representations are used to train a supervised model to predict the missing links. (5) Diffusion-based: These methods use techniques based on the analysis of the information gleaned from the movement of a random walker diffusion over the network (paths indicated by red arrows).

In this paper, we report the results of this community effort, where we evaluated the performance of various link prediction algorithms in the context of PPI prediction, and provided insights into the optimal computational tools required to detect the unmapped PPIs. In particular, we found that task-tailored similarity-based methods that leverage the underlying characteristics of PPIs show superior performance over other link prediction methods in both computational and experimental validations. We describe the details of these methods. The full datasets (including all of the benchmark interactomes and the experimental validation results), as well as the code for all of the tested methods and scoring functions, are freely provided to the scientific community.

## RESULTS

### 1) Computational Evaluation

Fig.3 summarizes the results of the computational evaluation of all tested methods using 10-fold cross validation. The metrics used to quantify the performance of each method are (i) AUROC: Area Under the Receiver Operating Characteristic; (ii) AUPRC: Area Under the Precision-Recall Curve; (iii) NDCG: Normalized Discounted Cumulative Gain; and (iv) P@500: Proportion of Positive PPIs, i.e., precision, in the top-500 prediction. To better rank the overall performance of different methods, we also computed a combined z-score for each method that summarizes its performance using different evaluation metrics. We next highlight the following key observations and insights from the analysis.

**Figure 3:**
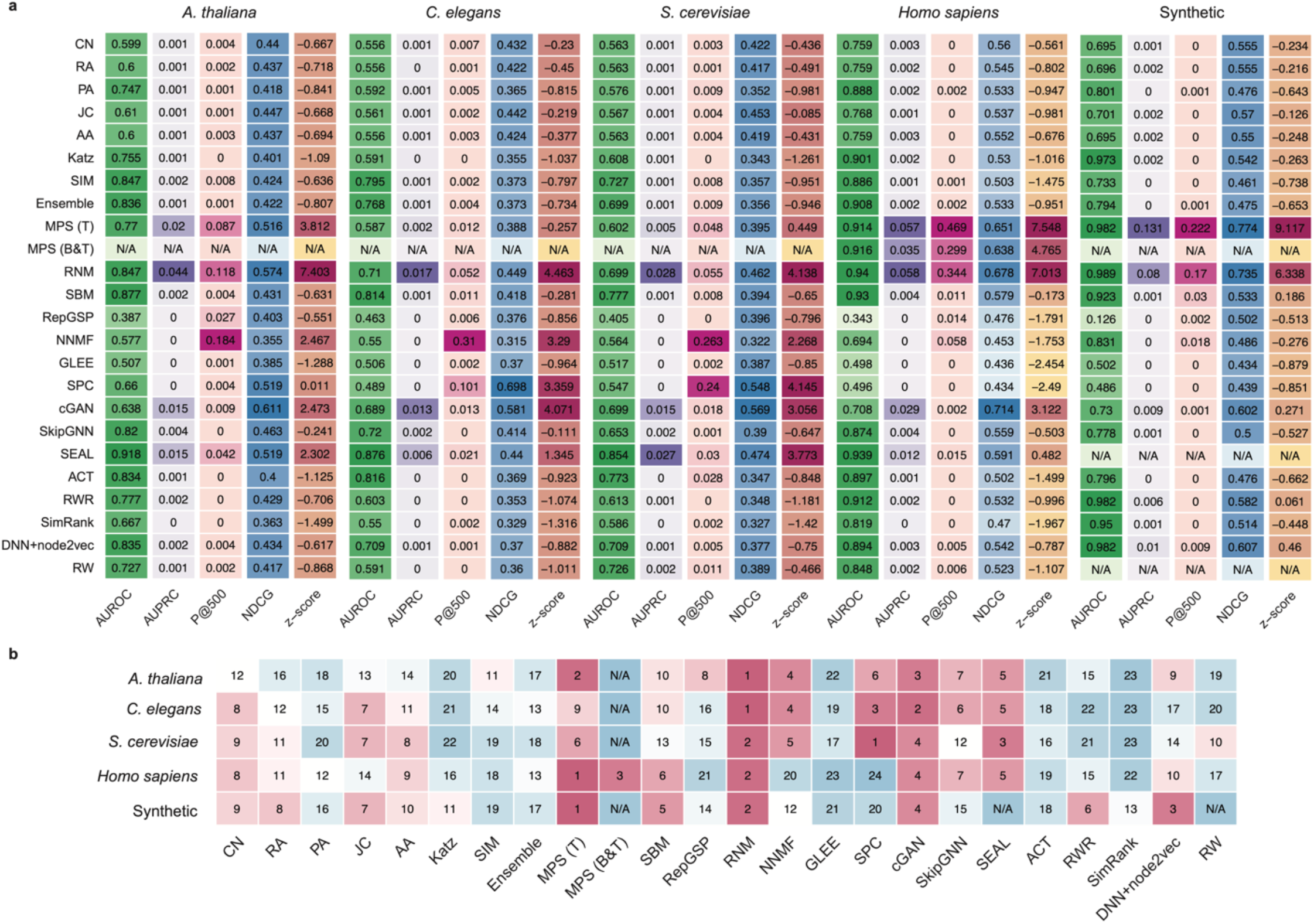
Computational evaluation of the PPI prediction methods. The details of each method are summarized in Table 1. **a**, Heatmap plots show the performance of each method on each interactome with the following evaluation metrics: Area Under the Receiver Operating Characteristic (AUROC), Area Under the Precision-Recall Curve (AUPRC), Precision of the top-500 predicted PPIs (P@500) and Normalized Discounted Cumulative Gain (NDCG). The overall performance is calculated from z-scores of three metrics. For each metric, darker color represents better performance. **b**, The ranking of the 24 methods on the five interactomes by z-scores. Note that, the performances of SEAL and RW in the synthetic interactome, and of MPS (B&T) in *A. thaliana*, *C. elegans*, *S. cerevisiae*, and synthetic interactomes are filled by N/A for consistency.

#### AUROC largely overestimates the performance of PPI prediction methods

Considering that the distribution of links is highly imbalanced in the PPI prediction problem due to the sparsity of interactome maps across organisms^19,20^, AUROC may overestimate the performance of a link prediction method, while AUPRC can provide more pertinent evaluation^21,22^. Indeed, by systematically comparing the performance metrics of various PPI prediction methods, we found clear evidence that AUROC largely overestimates the performance of any particular method. For example, the average AUROC over 10-fold cross validation of one of the top methods, SEAL, in *H. sapiens* is 0.94. This very high AUROC might lead us to mistakenly conclude that SEAL is an almost perfect prediction method since the maximum value of AUROC is 1. In fact, however, we found the average AUPRC of SEAL is only 0.012, implying a poor performance in finding PPIs (Fig.3a).

Since AUROC has been widely used in the link prediction literature^23–27^, we investigated whether or not the AUROC-based ranking of link prediction methods is consistent with rankings based on AUPRC or other metrics. First, we calculated the correlations between AUROC and other metrics over all methods. We found that AUROC is significantly correlated with AUPRC (Spearman *R* = 0.72, *p* < 2.2 × 10^−16^) and NDCG (Spearman *R* = 0.56, *p* = 1.6 × 10^−14^), but not with P@500 (Spearman *R* = 0.13, *p* = 0.14). Second, we found that overall the rankings of each method, including and excluding the AUROC metric, are quite consistent (Spearman *R* = 0.85, *p* < 2.2 × 10^−16^, SI Fig.S1). Note that the strong correlation between AUROC and AUPRC does not mean that the former is as good as the latter in evaluating the performance of any particular link prediction method. Instead, it just means we can still use AUROC to rank different methods (and the ranking will be roughly the same as if we use AUPRC), even though the AUROC values are systematically inflated (due to the data imbalance issue). Hereafter, we will, therefore, exclude AUROC in calculating the combined z-score for each method.

#### Predictability of interactomes is weak

Notably, both AUPRC and P@500 of most methods are quite small for all the five interactomes (Fig.3a-e). This observation suggests that successfully predicting missing links among a large unmapped PPI space remains a challenging task. To quantify the predictability of each interactome, we calculated its structural consistency index *σ*_*c*_ based on the first-order perturbation of the interactome’s adjacency matrix^28^. It is important to note that a network is highly predictable (with high *σ*_*c*_) if the removal or addition of a set of randomly selected links does not significantly change the network’s structural features (characterized by the eigenvectors of its adjacency matrix). We found that *σ*_*c*_ < 0.25 for all the five interactomes (SI, Fig.S2), which is much lower than that of social networks^28^ (e.g., Jazz^29^ and NetSci^30^ for which we have *σ*_*c*_ = 0.65 and 0.595, respectively). Such a low structural consistency indicates that the unobserved parts of the five interactomes considered in this project do not have very similar structural features as their currently observed part, which might be due to the high incompleteness of those interactome maps. (Indeed, for the social networks Jazz and NetSci, if we remove 90% of its links, their *σ*_*c*_ values drop from 0.65 to 0.16 ± 0.057 and from 0.595 to 0.25 ± 0.089, respectively (calculated from 50 random removals).

#### Performance of most PPI prediction methods vary considerably across different interactomes

We found that PPIs from *H. sapiens* and synthetic interactomes were predicted more accurately than PPIs from the other three organisms despite their very similar edge density (SI, Table S1). To better demonstrate the performance variability of the different methods, we ranked those methods based on their combined z-scores of AUPRC, AUROC, P@500 and NDCG. As shown in Fig.3b, some methods (e.g., RNM) show quite robust rankings (or consistent performance) across different interactomes, while the rankings of other methods show very large variability across interactomes. This variability is likely due, in part, to the different network characteristics, i.e., the number of links and average degree (SI, Table S1). To evaluate the performance variability systematically, we calculated the standard deviation of rankings across all five interactomes for each method, finding that RNM yielded the lowest variability. This suggests that RNM is robust and able to perform well on interactomes with quite different network characteristics (SI, Fig.S3).

#### Traditional similarity-based methods do not perform well

Traditional similarity-based link prediction methods have been heavily used in benchmark studies. These methods are based on a range of simple node similarity scores, such as the number of common neighbors, the Katz index, the Jaccard index, and the resource allocation index, to quantify the likelihood of potential links. Hence, they are more interpretable and scalable than other methods. However, the predefined local and global similarity score functions might significantly impact their predictive power for some networks, such as PPI networks. For example, the mere presence of two proteins in an interactome with many common neighbors does not necessarily imply that they should be connected to each other because interacting proteins are not necessarily similar and similar proteins do not necessarily interact^31^. We found that most of the traditional similarity-based methods, i.e., common neighbors (CN), Adamic-Adar index (AA), and Jaccard index (JC), yield negative z-scores, indicating their performance is below the average performance of all methods (Fig.3a).

#### There are three consistently high-performing methods

Among all the methods we tested, we found that RNM, MPS(T), and SEAL yield relatively high AUPRC and P@500 over all five interactomes. RNM is an ensemble method that integrates a diagonal noise model, a spectral noise model, and the L3 principle which shows that two proteins are expected to interact if they are linked by multiple paths of length three in the interactome^31^. RNM displays excellent performance across all five interactomes: it ranked No.1 in PPI prediction for *A. thaliana* and *C. elegans* interactomes, and No.2 for *S. cerevisiae, H. sapiens* and synthetic interactomes. MPS(T) also leverages the L3 principle, but at a higher level. In particular, it highly ranks a protein pair (*i*, *j*) if protein-*i* has similar neighborhood as that of protein-*j*’s neighbors. SEAL can leverage the sequence information of proteins. Moreover, it can learn a function that maps the subgraph patterns to link existence from a given network instead of using any traditional predefined similarity indices^32^. It is interesting to note that, compared with MPS(T), SEAL yields higher performance in the *C. elegans* and *S. cerevisiae* interactomes, but lower performance in the *H. sapiens* and synthetic interactomes. This performance difference might be due to the different graph mining techniques used by these methods. One of the similarity metrics in MPS(T) is defined as a function of the Jaccard index of the neighborhoods of two nodes, and it is reasonable that this metric should benefit from higher degree nodes for classification rather than those with small neighborhoods. Instead, SEAL uses Graph Neural Networks (GNNs) that define the features of a node by aggregating values from its nearest neighbors. In a case where nodes have many neighbors with similar features (e.g., nodes in the same biological modules), aggregation functions (e.g., the average function) could lead to nodes with similar embeddings, which are harder to distinguish.

#### Stacking models do not perform significantly better than individual methods within each interactome

It has been suggested previously that constructing a series of “stacked” models and combining them into a single predictive algorithm can achieve optimal or nearly optimal accuracy^33^. To confirm whether a stacking model is superior to individual methods in PPI prediction, we constructed four different stacking models. *Stacking-model-1* (Supervised): stack 36 individual topological predictors, which come in three types, global (functions of the entire network), pairwise (function of the joint topological properties of node pair *i*, *j*) and node-based (functions of the independent topological properties of node *i* and *j*), into a single algorithm, then train a classifier to predict the missing links. *Stacking-model-2* (Unsupervised): for each link, take the average of its ranking (in percentile form) calculated from RNM and MPS(T). *Stacking-model-3* (Unsupervised): for each link, take the maximum of its ranking calculated from RNM and MPS(T). *Stacking-model-4* (Unsupervised): for each link, we aggregated the rankings from RNM and MPS(T) using the Kemeny consensus^34^ approximated by the Dowdall^35^ or CRank^36^ method. Interestingly, we found that none of these stacking models can significantly outperform individual methods (see SI Fig.S4). In general, the advantage of Stacking-model-1 is that the meta-classifier can learn to select the best predictors through supervised training; thus, the overall performance of the stacking model outperforms any individual predictor. Here, the predictors of the two highest ranking methods can be directly used in PPI predictions without training a classifier. Moreover, the overlap between the PPIs predicted by different methods is very low (see SI Fig.S5 for Cohen’s kappa coefficient and the Jaccard index, and Fig.S8a for the overlapping of top-500 PPIs between the top-seven methods). Therefore, simply averaging the scores (Stacking-model-2) from different methods will decrease the ranking of those correctly predicted PPIs and, hence, degrade the predictive performance accordingly. Stacking-model-3 can yield slightly higher AUROC and NDCG than each individual method, but its AUPRC and P@500 still cannot surpass the best individual method. These results suggest that for a given network domain (e.g., PPI networks) stacking models do not always significantly outperform the best individual methods in link prediction.

#### Patterns of predicted PPIs

Based on their combined z-scores in the *H. sapiens* interactome, we selected the top-seven methods: MPS(T), RNM, MPS(B&T), cGAN, SEAL, SBM and DNN+node2vec. To examine whether there is any particular pattern among the top-500 PPIs predicted by each of the top-seven methods, we calculated the distribution of degree difference of proteins involved in the predicted PPIs (Fig.4a). We found that RNM, MPS(T), MPS(B&T), and SBM tend to predict PPIs involving proteins with larger degree difference than that of randomly selected PPIs in HuRI (with degree difference 16.56 ± 1.33 calculated from 10 random samplings of 500 PPIs in HuRI). The remaining methods cGAN, SEAL and DNN+node2vec tend to predict PPIs between nodes with similar degrees. In addition, for each method, we plotted the average degree of the proteins involved in their top-500 predicted PPIs on top of the degree distribution *P*(*k*) of the human interactome (with average degree 12.7) (Fig.4b). We found that RNM, SBM, MPS(T), and MPS(B&T) tend to predict PPIs involving proteins of high degrees, while the average degree of proteins in the top-500 PPIs from deep learning methods, such as DNN+node2vec and SEAL, is much lower. This difference could be due to the fact that RNM provides more predictions for high-degree nodes (see Fig.4b) and MPS also leverages the L3 principle at a high level, while DNN+node2vec focuses more on local network topological structure rather than on degree.

**Figure 4:**
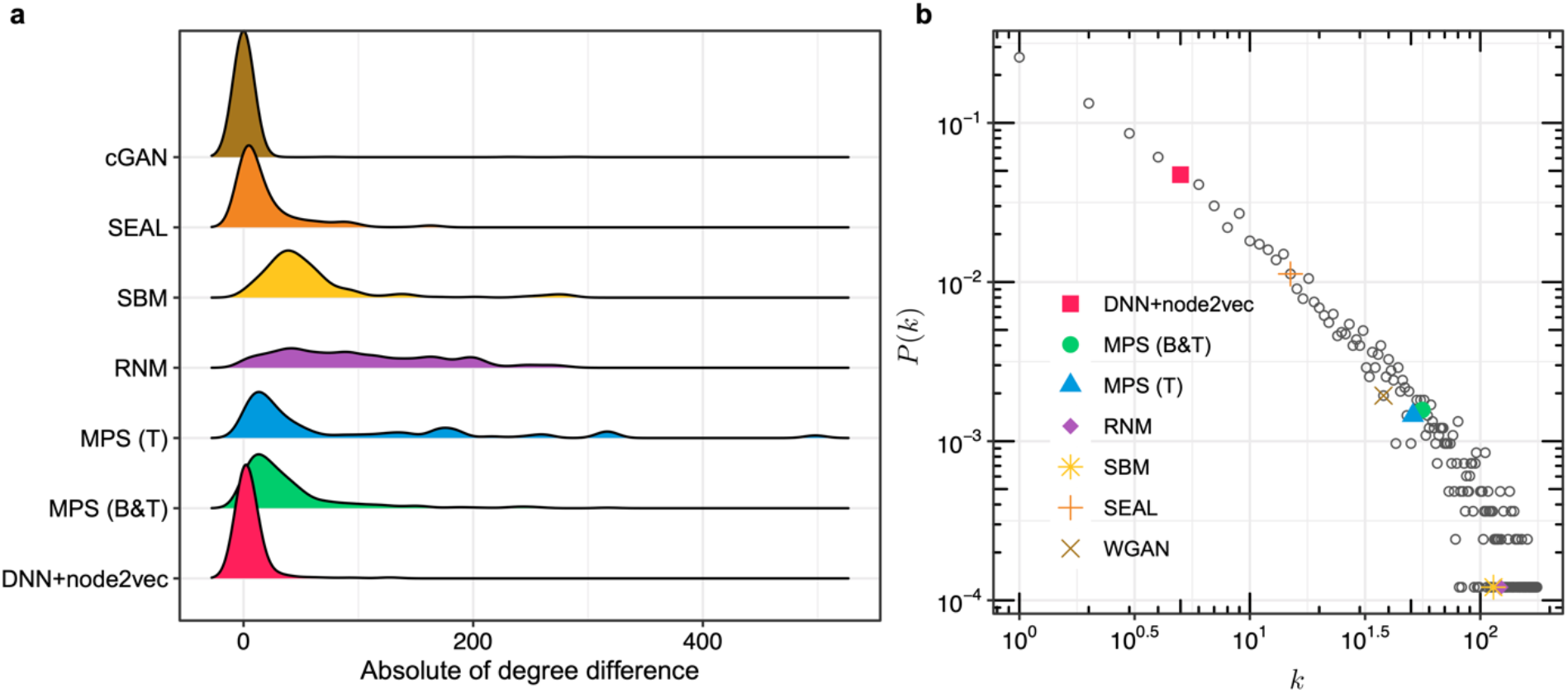
Patterns of top-500 PPIs predicted by the top-seven human PPI prediction methods. **a**, For these top-seven methods, we examined the distribution of absolute value between the degrees of each protein pair. **b**, Degree distribution of the *H. sapiens* interactome and the mean degree of proteins involved in the top-500 predicted PPIs of each method in log-log plot. *k* is the degree of a protein.

### 2) Experimental Validation

To the best of our knowledge, this is the first systematical benchmark study of network-based PPI prediction methods that incorporates large-scale confirmatory experimentation.

#### Performance of prediction methods in experimental validation

To validate the performance of the PPI prediction methods experimentally, we applied each of the top-seven methods (MPS(T), RNM, MPS(B&T), cGAN, SEAL, SBM and DNN+node2vec) to the human interactome (HuRI) and predicted the top-500 unmapped human PPIs. The union of the top-500 predicted human PPIs from top-seven methods includes 3,276 unique protein pairs. Next, we validated those protein pairs using the three orthogonal yeast two-hybrid (Y2H) assays formerly used to obtain the human interactome HuRI map. In total, we identified 1,177 new human PPIs (involving 633 proteins) by considering a protein pair to be *positive* if it is positive in at least one of the three assays, and *negative* if it is scored negative in all the three assays for which it was successfully tested. Overall, we found that MPS(B&T) is the most promising method in the sense that it simultaneously offers the highest number (376) of positive pairs and the lowest number (54) of negative pairs among its top-500 predicted PPIs (see Fig.5 and Table S3 for Precision).

**Figure 5:**
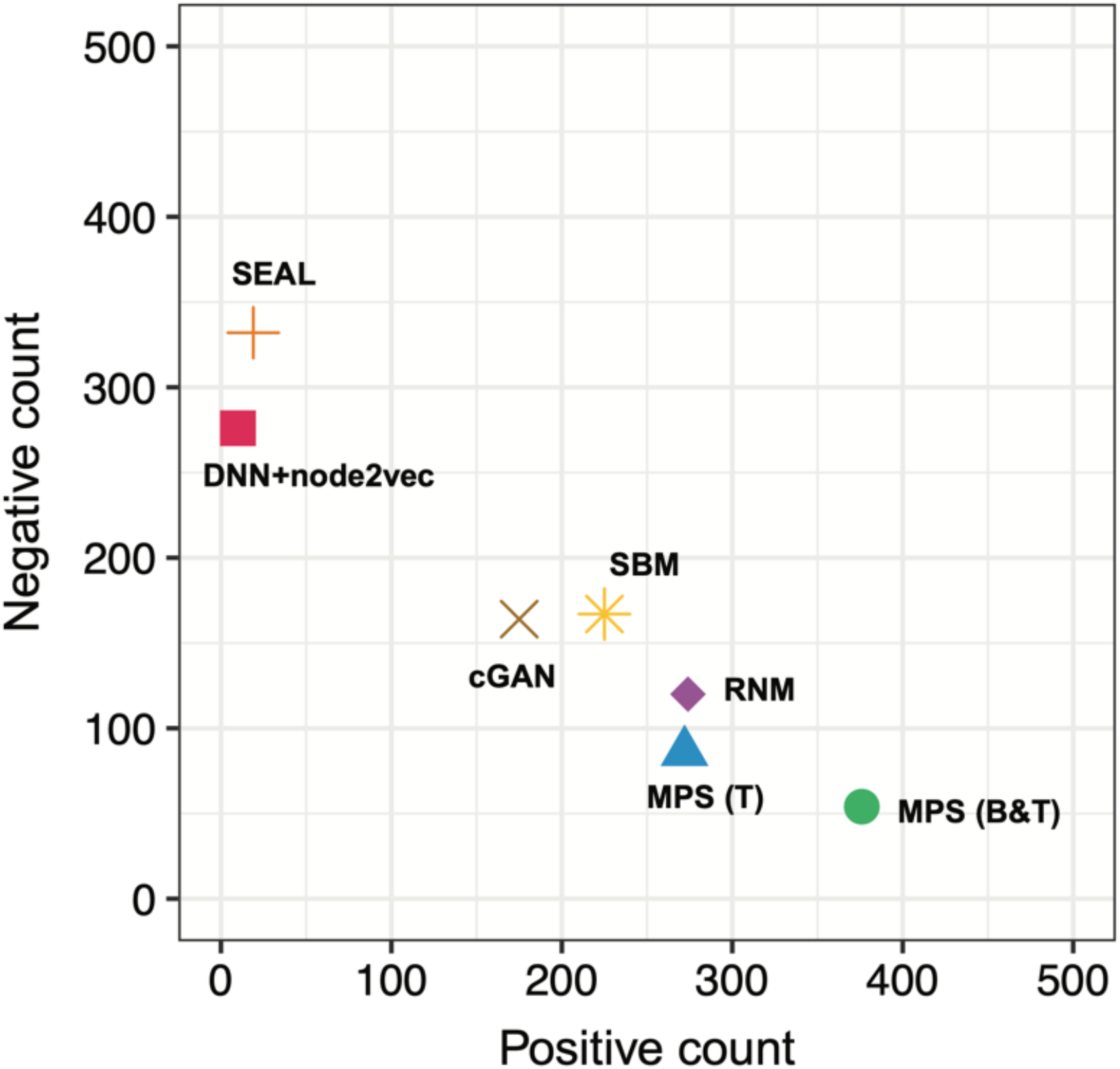
Experimental evaluation of the top-seven human PPI prediction methods. A protein pair is considered to be positive if it is positive in at least one of the three Y2H assays, and negative if it is negative in all the three assays. MPS(B&T) is the most promising method, which simultaneously offers the highest number (376) of positive pairs and the lowest number (54) of negative pairs among its top-500 predicted PPIs, yielding a precision of 87.4%. See Table S3 for the positive count, negative count, and the precision of other methods.

Interestingly, we found that most of the positive PPIs were uniquely predicted by a particular method (see Fig.S6 for the Venn diagram). Moreover, those PPIs simultaneously predicted by multiple methods tend to be positive. For example, the 11 PPIs simultaneously predicted by RNM, MPS(T) and MPS(B&T) are all positive.

Note that the human interactome map HuRI contains self-loops, i.e., some proteins interact with themselves, representing the diagonal elements of the adjacency matrix of HuRI. We understand that the prediction of diagonal elements is an orders of magnitudes easier task than the prediction of off-diagonal elements in the adjacency matrix, due to the much larger density of self-interactions: in HuRI, the average degree of those self-interacting proteins is 35.05, while the mean-degree of those non-self-interacting proteins is only 11.33. Among all prediction methods tested in this project, most of them tend to ignore self-loop prediction, but some of them (especially cGAN) do not. In fact, 495 of the top-500 PPIs predicted by cGAN are self-loops.

#### Combining predictions from the top three methods does not yield better precision

Those predicted PPIs with higher ranking positions (i.e., in the top of the top-500 list) presumably should have higher probabilities of being positive in experimental validation than those predicted PPIs with lower ranking (i.e., in the bottom of the top-500 list). To test this assumption, for each of the top-three methods in human PPI prediction, we plotted the ranking position distribution of the predicted PPIs that were validated to be positive in the Y2H experiments. As shown in Fig.S7a, surprisingly, these positive PPIs do not tend to appear more often at the top of the list. Instead, they appear almost randomly in the top-500 PPIs predicted by each method. (It is unclear if this intriguing phenomenon will continue to hold if we test more pairs, e.g., top-1000 PPIs.) Consequently, combining the top-500 PPIs predicted by those top-ranking methods into a new top-500 list does not yield a better performance in experimental validation. To demonstrate quantitatively this point, we combined the top-*N*_*k*_ PPIs predicted by MPS(B&T) and top-[(500 − *N*_*k*_)/2] PPIs from MPS(T) and RNM, respectively, with *N*_*k*_ ∈ [0,500] defined as a tuning parameter. We ensured that those PPIs predicted by different methods appear only once in the combined list. We found that the number of positive PPIs monotonically decreases with *N*_*k*_, indicating that combining the PPIs of greatest confidence predicted by different methods does not at all improve predictive performance (see SI Fig.S7b).

#### Structural and functional relationships of the validated new human PPIs

To explore the structural relationships of these predicted PPIs that were tested positive in the Y2H assay, we visualized the network constituted by them (in total 1,177 PPIs involving 633 proteins), finding four distinct clusters (Fig.6). These clusters were largely contributed by RNM, SBM, and MPS methods. We also found that the subnetworks contributed by RNM and SBM are close to each other. MPS(B&T) additionally contributes a cluster, besides the cluster formed with MPS(T) together. As shown in SI Table S2, those methods leveraging the connectivity features, i.e., MPS, RNM, and SBM, tend to predict PPIs in dense neighborhoods (with higher edge density and shorter characteristic path length) of the interactome. By contrast, deep learning methods (e.g., SEAL, and DNN+node2vec) tend to predict PPIs that are more scattered in the interactome, and the induced subgraphs have lower edge density and longer characteristic path length. To quantify the distance between the proteins involved in the positive PPIs predicted by different methods, we computed their network-based separation^3^ defined as *s*_*αβ*_ = 〈*d*_*αβ*_〉 − (〈*d*_*αα*_〉 + 〈*d*_*ββ*_〉)/2, where *α* and *β* represent the set of proteins involved in the positive PPIs predicted by two methods, respectively. 〈*d*_*αβ*_〉 is the average shortest distance between proteins in *α* and *β*, 〈*d*_*αα*_〉 (or 〈*d*_*ββ*_〉) is the average shortest distance between proteins within *α* (or *β*) in the original human interactiome^4^. We found that almost all methods predicted PPIs in specific and separated areas of the interactome (SI, Fig.S6b), as each method is more likely to reflect different topological characteristics.

**Fig.6.**
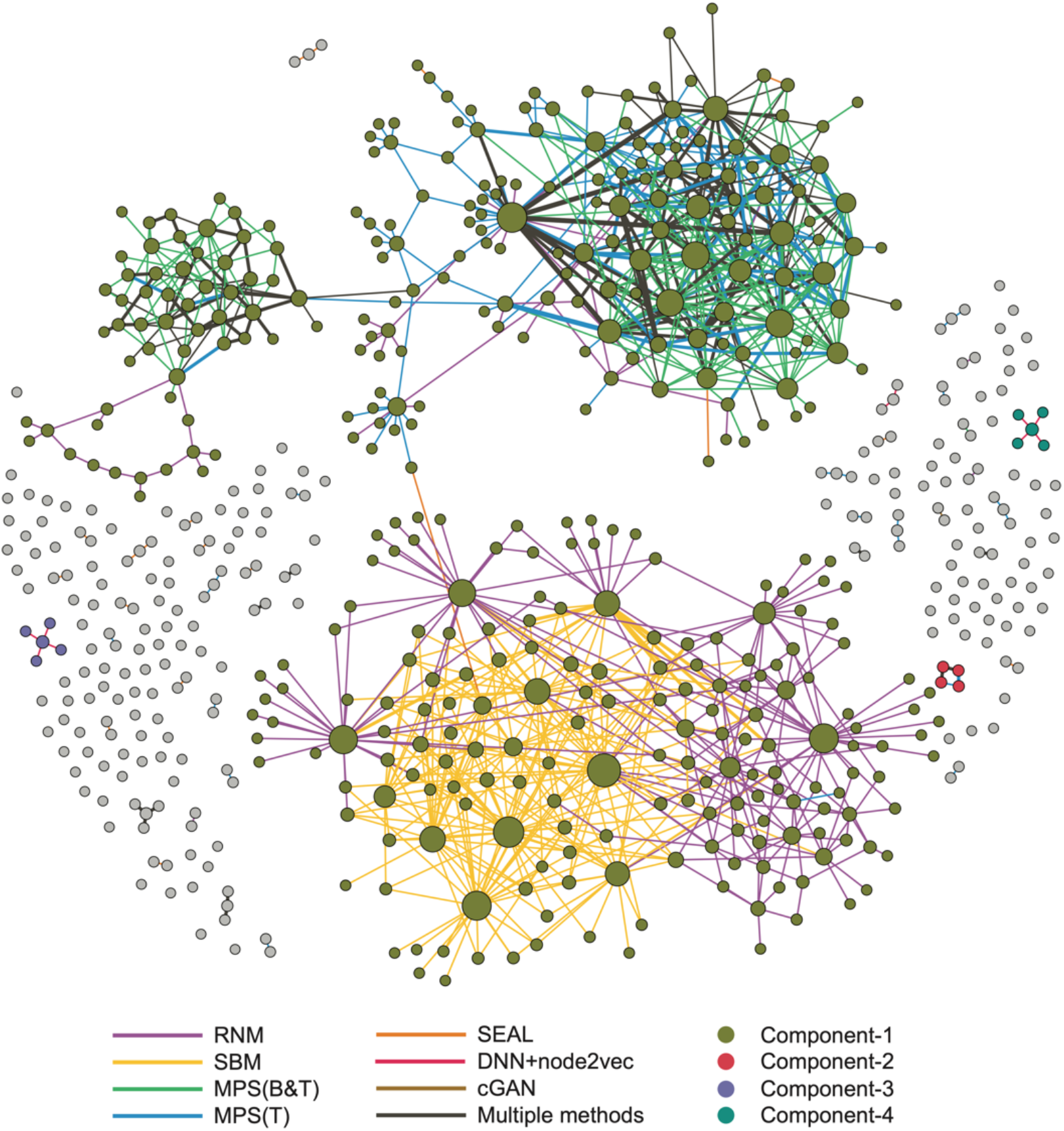
Structural relationships among new human PPIs. This network consists of all of the 1,177 new human PPIs predicted by the top-seven methods and validated by Y2H assays. Those PPIs that were predicted by a single method were colored based on the method that predicted them. Those PPIs that were predicted (i.e., among the top-500 predicted PPIs) by multiple methods were colored in black, with edge width proportional to the number of methods predicting this PPI. Nodes (proteins) are colored based on the connected component to which they belong. Node size is proportional to its degree. Note that there are in total 174 isolated nodes, representing self-interacting proteins (which were mostly detected by cGAN).

We also investigated the functional relationships of these positive PPIs, finding that they contribute to three functional modules, and each of those functional domains is associated with a distinctive, enriched GO term (SI, Fig.S8). This observation is also consistent with the previous finding that physical binding assembles proteins into large functional communities, thus providing insights into the global functional organization of the human cell^4^.

## DISCUSSION

As knowledge of human PPIs can help us understand complex biological and disease mechanisms, developing computational algorithms to discover previously unrecognized PPIs and, thereby, to improve the comprehensiveness of the human interactome map is critical. In order to achieve this goal, we have evaluated 24 representative network-based PPI prediction methods across five different interactomes using both computational and experimental validations.

As a result of this systematic evaluation and validation effort, we identified the top-performing methods that prove useful for PPI prediction. Our analysis showed that the predictive power of traditional similarity-based methods is limited, although they are more easily portable without the need to rely on organism-specific annotations. Furthermore, generic link prediction methods based on deep machine learning methods, including embedding and graph neural networks approaches, performed consistently across different interactomes studied in this project with a higher robustness, although their performances are not top-ranking. By contrast, link prediction methods specifically designed for PPI networks, e.g., MPS and RNM, displayed the most promising performance. More importantly, we found that different methods typically predicted positive PPIs that, rather than being scattered randomly in the interactome, are concentrated in specific areas (often associated with specific biological processes), and, furthermore, these areas overlap minimally among different methods. This minimal overlap is due to the underlying assumptions of each method that highlight particular network patterns, suggesting that we may need to use different methods simultaneously to reflect the variable patterns in the interactome and offer complementary predictions. From a network perspective, it would also be interesting to analyze in more detail the association between biological processes and the structural patterns they express in the interactome, a research topic that we leave for the future studies.

Conveniently, the top-ranking methods were robust and seemed suitable for all the interactomes we studied in this project. However, we cannot comment on the applicability to link prediction in general as we validated these methods only on PPI networks rather than networks from different scientific domains. In our analysis, stacking models did not show higher performance than any individual method in PPI prediction, which could be attributed to the low overlap between PPIs predicted by different method, as the overall search space is enormous. For consistency, in this project we focused on reference interactomes generated from the Y2H system and the experimental validations were also conducted using the Y2H system, which is one of the most popular and powerful tools to study PPIs. Of course, there are other PPI-mapping techniques available, e.g., mass spectrometry^37^. We anticipate that the top-ranking methods presented here will still offer excellent performance in predicting PPIs for interactomes mapped by other techniques.

In this project, although we tested many algorithms covering different categories, the classifiers based on protein sequence, for example, SVM^38–41^, RF^42,43^, FCTP^44^, and DPPI^45^, have not been tested for two reasons. First, those methods need to define a feature space for each link, which will lead to significant time complexity and memory requirement for HuRI (which has ~ 35 million unmapped PPIs). Second, structural information has relatively little impact on constructing the interactomes, primarily because there is a great difference between the number of proteins with known sequences and those with an experimentally determined three- (or four-) dimensional (i.e., tertiary or quaternary, respectively) structure^46^. In other words, this type of information is highly incomplete to be effectively exploited at the level of entire interactome. The recent success of AlphaFold^47^, a deep learning-based method to predict protein structure with atomic accuracy, is shedding light on resolving this limitation.

Despite their high performance compared to all 24 tested methods, the graph-mining-based deep learning strategies (i.e., SEAL, SkipGNN, and DNN+node2vec) are not in the top three rank order. A reason for this failing could be the difference in the patterns of predicted PPIs, as remarked upon previously. For example, DNN+node2vec tends to predict PPIs involving proteins with lower degree than the top three methods.

Based on these findings, we recommend the following considerations for effective PPI prediction. First, the method needs to leverage the inherent properties of the interactome to improve the predictive performance, such as avoidance of the use of first-order similarity between two proteins. Second, the unmapped PPI space is over several hundred times larger than the currently mapped space, causing a limited overlap of the most probable PPIs predicted by different methods, which obviously reduces the efficacy of ensemble or stacking models. Finally, incorporating protein sequence attributes into the network topology-based methods still requires the development of a (likely complicated) approach to combine them efficiently and effectively.

## MATERIALS and METHODS

### The INMC protein-protein interaction prediction project

This community effort was initiated by the International Network Medicine Consortium (INMC) aiming to provide a framework to assess the network-based computational methods in protein-protein interaction (PPI) prediction through standardized performance measures and common benchmarks. The INMC members were required to run their selected methods on five benchmark interactomes. For each interactome, members were required to submit two sets of results: (1) 10-fold cross-validation to compute the four performance measures; and (2) top-500 new human PPIs predicted by their methods by leveraging the whole human interactome. In total, we tested 24 link prediction methods. The top-500 new human PPIs provided by the top-7 high performance methods were further evaluated experimentally through yeast two-hybrid (Y2H) assays.

### Benchmark interactomes

To evaluate the performance of the considered computational methods, we need reliable and unbiased benchmark interactomes. Literature-curated interactomes of PPIs with multiple lines of supporting evidence might be highly reliable, but they are largely influenced by selection biases^48^. Therefore, here we focus on interactomes emerging from systematic screens that lack selection biases. For simplicity, we focus on binary datasets where co-complex membership annotations are not included. We used five interactomes for performance evaluation: (1) A plant interactome including 2,774 proteins and 6,205 PPIs, derived from the PPIs in the *A. thaliana* Interactome, version 1 (AI-1) and literature databases^15^; (2) a worm interactome including 2,528 proteins and 3,864 PPIs, derived from *C. elegans* version 8 (WI8), which is assembled from high-quality yeast two-hybrid PPIs^16^; (3) a yeast interactome of *S. cerevisiae* including 2,018 proteins and 2,930 PPIs, derived from the union of CCSB-YI1, Ito-core and Uetz-screen datasets^17^; (4) a human interactome including 8,274 proteins and 52,548 PPIs, derived from HuRI^4^, which is assembled from binary protein interactions from three separate high-quality yeast two-hybrid binding assays; and (5) a synthetic interactome including 8,274 proteins and 59,922 PPIs, generated by the duplication-mutation-complementation model^18^.

The duplication-mutation-complementation model constructs an interactome according to the following algorithm. To generate a graph *G*(*V*, *E*) with node set *V* and link set *E*, for each time step, a node is added to *G* according to one of two rules: duplication or divergence. In duplication, a node *i* is chosen at random and a new node *j* is connected to all the neighbors of *i*, while nodes *i* and *j* have a link with probability *p*. In divergence, the algorithm randomly chooses two nodes *i* and *j* and, for each of the nodes k linked to both nodes *i* and *j*, randomly selects one link, either (*i*, *k*) or (*j*, *k*), to be removed with probability *q*. The parameters *p* and *q* are chosen to fit key network properties of *G*, which we chose to be the human interactome. Here, we chose four network properties: mean degree, degree distribution (power-law exponent), clustering coefficient, and mean node betweenness. The degree distribution is estimated by maximum-likelihood fitting methods with goodness-of-fit tests based on the Kolmogorov-Smirnov statistic and likelihood ratios^49^. The optimal fitting is *p* = 0.125 and *q* = 0.435. The resulting interactome includes 8,274 proteins and 59,922 PPIs.

### PPI prediction methods

We compared in total 24 different methods that fall into five categories based on the adopted prediction strategy: similarity-based methods, probabilistic methods, factorization-based methods, machine learning methods, and diffusion-based methods (see Fig.2). Based on the information used in the prediction, these methods can also be divided into two categories: level-1 (based on network structure only) and level-2 (based on both network structure and node attributes) (see Table 1). Based on the usage of training PPIs labels, they can be further divided into supervised and unsupervised methods. In the following, we will provide an overview of each category. All the methods are briefly described in Table 1. Additional details can be found in Supplementary Information (SI).

- Similarity-based methods: these link prediction methods use a similarity score function based on local properties of network nodes to measure the likelihood of links. Two nodes with higher similarity score are considered to have a link between them with higher probability. For example, two nodes with more common neighbors are considered to be more similar and tend to link to each other. Note that when applied to PPI prediction, the adopted similarity measures are mostly based on interconnection properties of the nodes rather than on specific node features. The main advantages of these methods are that they are agnostic to any annotation-dependent features, making them easily applicable across organisms; and that they make few assumptions about global network structure.
- Probabilistic methods: The probabilistic and maximum likelihood algorithms assume that real networks have some structure, i.e., hierarchical or community structure. The goal of these algorithms is to select model parameters that can maximize the likelihood of the observed structure. As one of the most general network models, the stochastic block model (SBM) assumes that nodes are partitioned into groups with the probability that two nodes are connected depending solely on the groups to which they belong.
- Factorization-based methods: These methods use matrix factorization techniques to find a mapping to embed the original dimensional nodes in the network into a lower dimension so that similar nodes in the original network tend to have similar representation features. The resulting embedded lower-dimensional vectors (feature representations) can be used for many tasks, such as visualization, node classification, and link prediction. The link prediction task can be achieved by directly defining the likelihood of a link as the similarity of two nodes’ embedded features or using other complex classifiers, e.g., linear regression or deep neural networks.
- Machine learning: Machine learning (ML) is a growing field of pattern recognition algorithms that are trained on a given set of input data to make predictions based on the extracted patterns. Deep learning (DL) is a branch of machine learning composed of multi-layered neural network models. The recent success of deep neural networks is due to their ability to extract complex patterns in high-dimensional data by using non-linear functions. Graph neural networks (GNNs) are designed for learning over a graph. The graph convolution layers of the GNN are used to extract local substructure features for each node, and the graph aggregation layer aggregates node-level features into a graph-level feature vector^50^. These methods can learn parameters describing the general graph structural features and may include both node and connectivity features, showing promising performance in many network types^51^.
- Diffusion-based methods: These methods use techniques based on the analysis of the information gleaned from diffusion (typically from a random walker) over the network.

Details on methods using the above listed approaches can be found in the SI. We note that all surveyed methods use network (connectivity) information, while only a few (i.e., MPS(B&T), SEAL, RW2) incorporate information on protein information.

### Performance metrics

We assessed the performance of each protein interaction prediction method using four metrics: Area Under the Receiver Operating Characteristic (ROC) curve (AUROC), Area Under the Precision-Recall Curve (AUPRC), Precision of the top-500 predicted PPIs (P@500), and Normalized Discounted Cumulative Gain (NDCG). Notice that we included AUROC since it is widely used in the link prediction literature^23–27^ despite the fact that previous studies pointed out that AUROC is not a good performance metric for highly imbalanced data^21,22^. As the total number of PPIs for *C. elegans* and *S. cerevisiae* is less than 5,000, the test PPIs in 10-fold cross validation is less than 500, which means that the maximum P@500 is not 1. Given that the interactomes considered here are expected to be sparse, the number of true positive (i.e., existent) links will be dwarfed by the number of true negative (i.e., non-existent) links. In the literature, data imbalance is addressed by randomly selecting the same number of negative links to obtain a balanced validation list. Considering that we have also chosen three other metrics that can be used to quantify the classification methods and that are more robust to imbalanced data, we still reported the AUROC on the original imbalanced data.

Since the absolute values of AUROC and NDCG could be much larger than AUPRC and P@500, the values of AUPRC, P@500, and NDCG were each separately transformed into z-scores so that they have the same distribution, with mean value of 0 and standard deviation of 1. To compute a combined score that summarizes the performance of each method using the different evaluation metrics, we used the sum of the three z-scores, which is defined as:

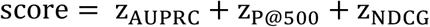

### Evaluation strategy

To validate the methods, we applied two strategies: computational validation and experimental validation. Computational validation refers to a computational assessment of the 24 methods using the aforementioned performance metrics. Experimental validation evaluates the top-seven methods (criteria for the selection of which are described below) according to their rank in the computational validation by applying wet laboratory experiments on their predicted PPIs.

### Computational validation

For each of the 24 methods, we performed computational validation using the 10-fold cross-validation approach. We randomly split the observed link set *E* into 10 subsets. For each iteration, one subset is selected as the probe set *E*^*P*^ and links in this subset are removed from the network. Links in the remaining nine subsets constitute the training set *E*^*T*^ and form the residual network. Note that some methods model both existing and non-existing links in training. For these methods, we added to the training set negative (non-existing) links generated by using balanced random sampling^52,53^. To compare different methods in a systematic way, we computed the aforementioned performance metrics considering the test set as the union of the probe set *E*^*P*^ and all the non-existing links not in the training set.

### Selection of the top-seven methods in human PPI prediction

Based on the results of the computational validation, we selected the top-seven methods in human PPI prediction (based on their combined z-scores) for further experimental validation. For each of the top-seven methods, we validated its top-500 predicted PPIs based on the whole human interactome. We found that the combined z-scores of RNM, MPS(T), MPS(B&T), cGAN, SEAL, and SBM are much higher than all other methods, suggesting their superior performance over other methods. Additionally, we chose DNN+node2vec as the seventh method to be validated experimentally, because its P@500 is higher than that of the next three best-performing methods, SkipGNN, AA, and CN.

### Experimental validation

The union of the top-500 human PPIs predicted by the top-seven methods includes 3,276 unique protein pairs. We systematically tested those protein pairs by performing three complementary yeast two-hybrid (Y2H) assays as performed previously for the human interactome^4^. Briefly, 5 μl of glycerol stocks of Y8930:DB-ORF and Y8800:AD-ORF haploid strains were cherry picked from the hORFeome collection into 200 μl of selective media (Synthetic Complete media without Leucine [SC-Leu] or Synthetic Complete media without Tryptophan [SC-Trp], respectively) and arrayed to generate the pairs to be tested. After overnight growth, 5 μl of Y8930:DB-ORF and 5 μl of Y8800:AD-ORF culture were transferred into YEPD (Yeast Extract Peptone Dextrose). After incubating overnight at 30 °C to allow mating to occur, 10 μl of yeast culture were transferred into 120 μl SC-Leu-Trp media to allow for selection of diploid yeast cells. The next day, diploid yeast cultures were spotted on Synthetic Complete media without leucine, tryptophan and histidine with 1 mM 3-Amino-1,2,4-triazole (SC-Leu-Trp-His+1 mM 3AT) to test for interactions and SC-Leu-His+1 mM 3AT supplemented with either 1 mg/L cycloheximide (CHX) for assay version 1 or 10 mg/L for assay versions 2 and 3 to test for auto-activation. After 72 h incubation at 30 °C, diploid cells that grew on SC-Leu-Trp-His+3AT media but not on SC-Leu-His+3AT+CHX media were scored positive. If a pair had a similar or higher frequency of yeast colonies on the CHX plate compared to the retest plate, then it was scored as a spontaneous auto-activator (AA). Cases showing contamination or where no diploid yeast were spotted (for example due to a pipetting failure), were reported as “not-tested.” To benchmark the performance of the experiment a set of positive controls and a set of random controls were included in each test. Experiments were only considered complete where both controls performed as expected. Details are described in Ref.[4] and http://www.interactome-atlas.org.

## Data and code availability

All data and scripts are available at https://github.com/spxuw/PPI-Prediction-Project.

## Acknowledgement

L.M., A.F. and L.B. were partially supported by the ERC Advanced Grant 788893 AMDROMA “Algorithmic and Mechanism Design Research in Online Markets”, the EC H2020RIA project “SoBigData++” (871042), and the MIUR PRIN project ALGADIMAR “Algorithms, Games, and Digital Markets”. F.L. was supported by a Wallonia-Brussels International (WBI)-World Excellence Fellowship. P.F. and B.Á. were supported by the National Research, Development and Innovation Office of Hungary (Research Excellence Program - TKP, National Heart Program NVKP 16-1-2016-0017) and by the Higher Education Institutional Excellence Program of the Ministry of Human Capacities in Hungary, within the framework of the Therapeutic Development thematic program of the Semmelweis University. Y.-Y.L. acknowledges grants from National Institutes of Health (R01AI141529, R01HD093761, RF1AG067744, UH3OD023268, U19AI095219, and U01HL089856). JL acknowledges support from the National Institutes of Health (R01 HL155107, R01 HL155096, U01 HG007690, and U54 HL119145); and from the American Heart Association (D700382 and CV-19).

## Author contributions

Y.-Y.L. and P.V. conceived and designed the project. L.M., A.F. and L.B. developed and tested the MPS(T) and MPS(B&T) methods. T.W. and I.K. developed and tested the RNM method. O.M.B., B.B., M.P., B.Á. and P.F. developed and tested the cGAN method. L.V. and J.M. developed and tested the DNN+node2vec method. S.C., M.P., G.S., and F.C. developed and tested the RepGSP method. L.M. tested the RW method. X.-W.W. tested the other 17 methods. K.S., T.H., F.L., L.W., J.-C.T. and M.C. conducted the experimental validations. X.-W.W., L.M., P.V. and Y.-Y.L. analyzed the results. X.-W.W. and Y.-Y.L. wrote the manuscript with assistance from L.M., P.V., and K.S.. L.B., I.K., B.Á., L.V., S.C., M.A.C., A.-L.B., E.K.S., and J.L. edited the manuscript. All authors approved the final manuscript.

