## Supplementary Information for "Assessment of community efforts to advance computational prediction of protein-protein interactions"

Xu-Wen Wang[1], Lorenzo Madeddu[2], Kerstin Spirohn[3,4,5], Leonardo Martini[6], Adriano Fazzone[6], Luca Becchetti[6], Thomas P. Wytock[7], István A. Kovács[7,8], Olivér M. Balogh[9], Bettina Benczik[9,10], Mátyás Pétervári[9], Bence Ágg[9,10], Péter Ferdinandy[9,10] , Loan Vulliard[11,12], Jörg Menche[11,12,13], Stefania Colonnese[14], Manuela Petti[6], Gaetano Scarano[14], Francesca Cuomo[14], Tong Hao[3,4,5], Florent Laval[3,4,5,15,16,17], Luc Willems[15,17], Jean-Claude Twizere[16,17], Michael A. Calderwood[3,4,5], Enrico Petrillo[18,19], Albert-László Barabási[18,20,21], Edwin K. Silverman[1], Joseph Loscalzo[18], Paola Velardi[2], Yang-Yu Liu[1]

[1]*Channing Division of Network Medicine, Department of Medicine, Brigham and Women's Hospital and Harvard Medical School, Boston, MA, 02115, USA.*

[2]*Translational and Precision Medicine Department Sapienza University of Rome, Rome, Italy.*

[3]*Department of Computer, Control, and Management Engineering "Antonio Ruberti", Sapienza University of Rome.*

[4]*Department of Physics and Astronomy, Northwestern University, Evanston, IL 60208-3112, USA.*

[5]*Northwestern Institute on Complex Systems, Northwestern University, Evanston, IL 60208.*

[6]*Cardiometabolic and MTA-SE System Pharmacology Research Group, Department of Pharmacology and Pharmacotherapy, Semmelweis University, Budapest, Hungary.*

[7]*Pharmahungary Group, 6722 Szeged, Hungary.*

[8]*CeMM Research Center for Molecular Medicine of the Austrian Academy of Sciences, Vienna, Austria.*

[9]*Department of Structural and Computational Biology, Max Perutz Labs, University of Vienna, Vienna, Austria.*

[10]*Faculty of Mathematics, University of Vienna, Vienna, Austria.*

[11]*Department of Information Engineering, Electronics, and Telecommunications (DIET), University of Rome "Sapienza", Rome, Italy.*

[12]*Center for Cancer Systems Biology (CCSB), Dana-Farber Cancer Institute, Boston, MA, USA.*

[13]*Department of Genetics, Blavatnik Institute, Harvard Medical School, Boston, MA, USA.*

[14]*Department of Cancer Biology, Dana-Farber Cancer Institute, Boston, MA, USA.*

[15]*Laboratory of Molecular and Cellular Epigenetics, GIGA Institute, University of Liège, Liège, Belgium.*

[16]*Laboratory of Viral Interactomes, GIGA Institute, University of Liège, Liège, Belgium.*

[17]*TERRA Teaching and Research Centre, University of Liège, Gembloux, Belgium.*

[18]*Department of Medicine, Brigham and Women's Hospital and Harvard Medical School, Boston, MA, USA.*

[19]*Department of General Internal Medicine and Primary Care, Brigham and Women's Hospital, Boston, MA, USA.*

[20]*Network Science Institute and Department of Physics, Northeastern University, Boston, MA, USA.*

[21]*Department of Network and Data Science, Central European University, Budapest H-1051, Hungary.*

(Dated: September 22, 2021)

### CONTENTS

I. Categories of PPI prediction methods 4
  A. Similarity-based methods 5
  B. Probabilistic methods 12
  C. Factorization-based methods 14
  D. Diffusion-based methods 15
  E. Machine Learning method 17

II. Supplementary tables 21

III. Supplementary figures 25

References 33

#### I. CATEGORIES OF PPI PREDICTION METHODS

Protein-protein interaction (PPI) networks, or the interactome, describe physical interactions between proteins that mediate signaling, regulatory, and transport events in the cell, or assemble into (multi-) protein complexes [1–3]. Interactomes are a specific category of complex network [4–7]. Therefore, link prediction applicable to general complex networks can also be directly applied. In the following, we describe the existing link prediction methods for general networks and some methods specifically designed for PPI prediction.

A network can be mathematically represented by a graph $G(\mathcal{V}, \mathcal{E})$, where $\mathcal{V} = \{1, 2, \ldots, N\}$ is the node set and $\mathcal{E} \subseteq \mathcal{V} \times \mathcal{V}$ is the link set. A link is a node pair $(i, j)$ with $i, j \in \mathcal{V}$, representing a certain interaction, association or physical connection between nodes $i$ and $j$. Link prediction aims to infer the missing links or predict future links between currently unconnected nodes based on the observed links [8–10].

In this survey, we consider network-based methods, that is, methods that mostly or exclusively leverage local or global connectivity properties of interactomes, rather than biological features, such as protein sequence or three-dimensional structure [11]. One advantage of connectivity-based methods is that some biological features like three-dimensional structure are currently known only for a small percentage of proteins in the interactome, thereby limiting their usage in large-scale PPI prediction, which is the aim of this paper. We grouped the prediction strategies considered in this study, into five classes: similarity-based methods, probabilistic methods, factorization-based methods, diffusion-based methods, and machine learning methods. Similarity methods, traditionally used in the network link prediction literature, exploit the similarity of connectivity patterns of individual nodes in the network. Probabilistic methods define a probability distribution over unobserved links. Factorization-based methods attempt to capture the information of both global structure and clustering structure of a network, using low rank approximations as well as blocks in networks' adjacency matrices. Diffusion methods are stochastic processes, such as random walks, that can be used to extract information about nodes or dense groups of nodes in a network, and to predict interactions. Finally, researchers have explored several machine learning techniques ranging from classic learning methods to deep learning methodologies. Popular deep learning approaches for link prediction rely on novel techniques such as the Graph Neural Networks (GNNs), to exploit complex patterns in network topology.

##### A. Similarity-based methods

In similarity-based methods, each non-observed node pair $(i, j)$ is assigned a similarity score $s_{ij}$. A higher score is assumed to represent a greater probability of link existence. The similarity score can be defined in many different ways and may rely on local information only, or some combination of local and global information

**Common Neighbors**. The common neighbors algorithm quantifies the overlap or similarity of two nodes as follows [12]:

$$s_{ij} = |\Gamma(i) \cap \Gamma(j)|, \tag{1}$$

where $\Gamma(i)$ denotes the set of neighbors of node $i$, $\cap$ denotes the intersection of two sets and $|X|$ denotes the cardinality or size of set $X$.

**Adamic-Adar Index**. This similarity index measures the similarity between two entities based on the connectivity of their shared neighbors [13]:

$$s_{ij} = \sum_{m \in \Gamma(i) \cap \Gamma(j)} \frac{1}{\log|\Gamma(m)|}. \tag{2}$$

**Jaccard Index**. The Jaccard index measures the overlap of two nodes normalized by their total number of neighbors [14]:

$$s_{ij} = \frac{|\Gamma(i) \cap \Gamma(j)|}{|\Gamma(i) \cup \Gamma(j)|}, \tag{3}$$

where $\cup$ denotes the union of two sets.

**Cosine Similarity (i.e. Salton Index)**. The cosine similarity is closely related to the Jaccard index, with the normalization as the product of the node degrees:

$$s_{ij} = \frac{|\Gamma(i) \cap \Gamma(j)|}{\sqrt{|\Gamma(i)||\Gamma(j)|}}. \tag{4}$$

**Preferential Attachment Index**. This index assumes that the existent likelihood of a link between two nodes is proportional to the product of their degrees [15]:

$$s_{ij} = k_i k_j, \tag{5}$$

**Resource Allocation Index**. This index is based on a resource allocation process between pairs of nodes [16, 17]. The similarity between a node pair $(i, j)$ is defined as the amount of resource $j$ received from $i$ through their common neighbors:

$$s_{ij} = \sum_{m \in \Gamma(i) \cap \Gamma(j)} \frac{1}{k_m}. \tag{6}$$

Here, we assume each common neighbor has a unit of resource and will equally distribute among all its neighbors.

**Hub Promoted Index**. This index is used to measure the link formation between hub nodes and low-degree nodes [18]:

$$s_{ij} = \frac{|\Gamma(i) \cap \Gamma(j)|}{\min(|\Gamma(i)|, |\Gamma(j)|)}. \tag{7}$$

**Hub Depressed Index**. This index is used to measure the link formation between hub nodes [18]:

$$s_{ij} = \frac{|\Gamma(i) \cap \Gamma(j)|}{\max(|\Gamma(i)|, |\Gamma(j)|)}. \tag{8}$$

**Sorensen Index**. This index is very similar to Jaccard index but with more robustness against the outliers [19]:

$$s_{ij} = \frac{|\Gamma(i) \cap \Gamma(j)|}{k_i + k_j}. \tag{9}$$

**L3 Index**. This index is based on the network paths of length three [20]:

$$s_{ij} = \sum_{u,v} \frac{a_{iu} a_{uv} a_{vj}}{\sqrt{k_u k_v}}, \tag{10}$$

where $a_{iu} = 1$ if node $i$ and $u$ interact, and zero otherwise.

**Sim Index**. This index combines the Jaccard index and the L3 index together [21]:

$$S = AJ + JA, \tag{11}$$

where $A$ is the adjacency matrix and $J$ is the Jaccard similarity matrix.

**Katz Index**. The Katz index is based on a weighted sum over the collection of all paths connecting nodes $i$ and $j$ [18]:

$$s_{ij} = \sum_{l=1}^{\infty} \beta^l (A^l)_{ij}, \tag{12}$$

where $\beta$ is a damping factor that gives the shorter paths more weights, and $A$ is the adjacency matrix of the network. The $N \times N$ similarity matrix $S = (s_{ij})$ can be written in compact form as [22]:

$$S = (I - \beta A)^{-1} - I, \tag{13}$$

where $I$ is the identity matrix. The damping factor $\beta$ is a free parameter and should be less than the reciprocal of the absolute value of the largest eigenvalue $|\lambda_{\max}|$ of $A$.

**Structural Perturbation Method**. This method assumes that a group of links is predictable if removing them has only a small effect on the eigenvalues of the network's adjacency matrix, which serve as proxies for the network's more general structural features [23]. This method proceeds by dividing an existing network $A$ into a "training" subset that is used to calculate the eigenvalues and eigenvectors of the remaining network $A^R_{ij} = \sum_{k=1}^{N} x_{ik} x_{kj} \lambda_k$, and a "validation" subset that will be used to characterize the predictability, $\Delta E$, which has size $L$ [23]. The $\Delta E$ is projected onto the eigenvectors, $\Delta\lambda_k = \frac{\sum_l \sum_m x_{kl} \Delta E_{lm} x_{mk}}{\sum_l x_{kl} x_{lk}}$, where $\Delta\lambda_k$ is the shift in eigenvalues that best approximates $\Delta E$. The best-fit adjustments are added to the eigenvalues and multiplied by the eigenvectors to obtain a perturbed adjacency matrix: $\tilde{A}_{ij} = \sum_{k=1}^{N} (\lambda_k + \Delta\lambda_k) x_{ik} x_{kj}$. The $L$ largest elements of $\tilde{A}_{ij}$ are selected, such that $A_{ij} \neq 1$, resulting in a set of edges called $E^L$. The Structural Perturbation Index is then $|E^L \cap \Delta E|/L$.

When applying the structural perturbation method to a given network, a byproduct is the so-called structural consistency index:

$$\sigma_c = \frac{|E^L \cap \Delta E|}{|\Delta E|}, \tag{14}$$

which can be used to quantify the predictability of the network.

**MPS**. This method is based on three measures: MaxSimScore, PAscore, and SeqScore [8, 21, 24, 25]. More formally, given two proteins $u$ and $v$, their Jaccard Index $J(u,v)$ is defined as:

$$J(u,v) = \frac{|\Gamma(u) \cap \Gamma(v)|}{|\Gamma(u) \cup \Gamma(v)|}$$

where $\Gamma(u)$ is the set of $u$'s neighbors in the PPI network. The Jaccard Index can be used to identify proteins with similar interfaces, and we can leverage this information to score potential protein interactions.

To better understand how Jaccard Index can be useful in scoring potentially interacting pairs, consider the following scenario: assume we have a protein pair $(u,v)$ for which we want to assess likelihood of interaction. If $u$ is similar to the neighbors of $v$ (i.e., they largely share the same interaction partners) and $v$ is similar to the neighbors of $u$, then it is more likely for $u$ and $v$ to share complementary binding sites. Thus, they are more likely to interact.

More formally, given proteins $u$ and $v$, we define their *interaction likelihood* as follows:

$$\text{MaxSimScore}_{\text{Topological}}(u,v) = \max_{x \in \Gamma(v)} J(u,x) + \max_{y \in \Gamma(u)} J(y,v),$$

where $\Gamma(u)$ is the set of neighbors of $u$. The proposed method is a modification of the `sim` method proposed by Chen et al [21], where the sums are replaced by max functions.

Assuming that proteins that are biologically similar to other interacting proteins are more likely to interact, we designed a framework that leverages proteins' primary sequence and priori knowledge of interacting proteins to identify new candidate interactions. This framework is heavily inspired by the PIPE algorithm [24], a well known protein interaction prediction engine that leverages recurring short polypeptide sequences between known interacting protein pairs to score candidate pairs. In other words, given a candidate pair $(u,v)$ and a known interacting pair $(a,b)$, if multiple regions $u_i$ of $u$'s primary structure are similar to analogous regions in $a$ and the same happens for $v$'s primary structure resemble with respect to $b$, $u$ and $v$ are more likely to be a interacting protein pair.

Starting from PIPE [24], we designed a simplified framework in which we don't directly consider all co-occurrences of two sub sequence pairs $(u_i, v_j)$, but we use them to define a protein similarity measure. In more detail, we define the biological similarity of two proteins $u$ and $a$ as follows:

$$S(u,a) = \begin{cases} 1, & \text{if } J'(s_u, s_a) > 0 \\ 0, & \text{otherwise} \end{cases} \tag{15}$$

where $J'(s_u, s_a)$ is the Jaccard Index between the set of shingles of the primary structures of protein $u$ and $a$. Given a pair of no-interacting proteins $(u, v)$ and the knowledge graph of interacting proteins $G(V, E)$, the algorithm computes the likelihood of the interaction as the following steps:

- **Step 1:** generate $A$, the set of proteins that are biologically similar to $u$. More formally, $A$ is defined as follows:

$$A = \left\{ a \in V \middle| S(u,a) = 1 \right\} \tag{16}$$

- **Step 2:** generate $R$, the set of proteins that interact with at least one protein in $A$. We can define $R$ as follows:

$$R = \left\{ x \in V \middle| \exists a \in A | (x,a) \in E \right\} \tag{17}$$

- **Step 3:** generate $B$, the set of proteins that are biologically similar to $v$. This set is defined the same way we defined $A$:

$$B = \left\{ b \in V \middle| S(v,b) = 1 \right\} \tag{18}$$

- **Step 4:** compute the likelihood of the interaction $(u, v)$ as the size of intersection between $R$ and $B$( i.e. $\text{DirSimScore}_{\text{Biological}}(u, v) = |R \cap B|$).

- **Step 5:** compute $\text{DirSimScore}_{\text{Biological}}(v, u)$ by executing, another time, steps from 1 through 4, this time switching $u$ with $v$.

- **Step 6:** return the following score as a measure of the interaction likelihood of proteins $u$ and $v$:

$$\text{MaxSimScore}_{\text{Biological}}(u,v) = \max\left(\text{DirSimScore}_{\text{Biological}}(u,v), \text{DirSimScore}_{\text{Biological}}(v,u)\right) \tag{19}$$

For each protein $u$, our framework pre-computes the set $A$ with a computational cost of $O(P^2L)$ where $P$ is the number of proteins and $L$ is the average protein primary sequance length. This way, steps 1 and 2 have constant time complexity. Finally, it finds $R$ in and computes the size of the intersection in $O(P)$. Thus, the overall time complexity is $O(P^2L)$ that is very small if we compare it with the time complexity of PIPE that is $O(P^3L^2)$.

Finally, we create a framework that leverages both biological and topological information, by aggregating the topological and the biological scores of a candidate pair $(u, v)$ via a convex combination. More formally, we defined the final likelihood of a candidate pair of proteins $(u, v)$ as:

$$\text{MPS(T \& B)}_\beta(u, v) = \beta \cdot \text{MaxSimScore}_{\text{Topological}}(u, v) + (1 - \beta) \cdot \text{MaxSimScore}_{\text{Biological}}(u, v) \tag{20}$$

where $\beta \in \big[0, 1\big]$, and with both scores suitably normalized before the computation of the sum.

We refer to the $\text{MPS(T \& B)}_\beta(u, v)$ method as the **MPS(T)** method, when $\beta = 1$; in a similar fashion, when $\beta = 0.5$, we refer to the $\text{MPS(T \& B)}_\beta(u, v)$ method as the **MPS(T & B)** method. The code for MPS is available at: `https://github.com/spxuw/PPI-Prediction-Project`.

**RNM**. This method is an extension of the $L3$ method, which has previously been shown to be effective in predicting PPIs [20, 26]. The $L3$ method has no free parameters, and thus makes predictions irrespectively of the amount of incompleteness in the data. In practice, we need less predictions when the measured network is almost complete and more if the dataset is missing most connections. We expect that a method with at least one adjustable parameter can account for the amount of incompleteness and should be able to out-perform the standard $L3$ method.

Our starting point is an edge prediction method of the form

$$\mathbf{P} = \mathbf{X}\mathbf{A}\mathbf{X}^\intercal \tag{21}$$

where $A$ is the $n \times n$ adjacency matrix of the observed PPI network, $\mathbf{X}$ is an $n \times n$ matrix to be determined, and $n$ is the number of proteins in the screen. In the case of $L3$, $\mathbf{X}_{\text{L3}} = \mathbf{A}\mathbf{D}^{-1/2}$, where $\mathbf{D} = \text{diag}(d_i)$, the matrix with the degrees of the nodes of $\mathbf{A}$ on the diagonal. In addition, we consider two alternative methods of the form $\mathbf{X} = |\mathbf{A}|(|\mathbf{A}| + \mathbf{N})^{-1}$, with

$|A| = \sqrt{A^2}$ and $\mathbf{N}_1 = \mathrm{diag}\left(a d_i^{1/c}\right)$ in the first method. In the second method, we first decompose $\mathbf{A} = \mathbf{U}\boldsymbol{\Lambda}\mathbf{U}^\intercal$, where $\boldsymbol{\Lambda} = \mathrm{diag}\lambda_i$ with $\lambda_1$ being the largest eigenvalue. Then, we consider a method of the form $\mathbf{N}_2 = \mathbf{U}\,\mathrm{diag}(M)\,\mathbf{U}^\intercal$, where $M = b\lambda_1$. Our predictions were made with the parameters $a = 3$, $c = 3$, and $b = 0.95$. Both methods give a small number of new predictions when $N$ is negligible compared to $|A|$, i.e. when $a$ or $b$ is sufficiently small.

Not knowing *a priori* which method is expected to perform best, we choose to use an average of the predictions of the three methods. For the internal cross-validation, we define $r_{ij}$ to be the normalized rank $i^{\text{th}}$ interaction predicted by the $j^{\text{th}}$ method, with the highest score being $r_{ij} = 1$. Then, we use $\bar{r} = \sum_j r_{ij}/3$ to rank the predictions and evaluate against the test set in terms of the required statistics.

For the external validation, predictions are made for all three methods for each of 3 assays in the human interactome. Within each assay, the predictions are merged and averaged as described in the preceding paragraph, except that the ranks are unnormalized. Since the assays cover largely disjoint sets of genes, the absence of a prediction of a particular pair in one assay can result from that assay's failure to cover the pair in question, or it may reflect a genuine absence of interaction. To merge the predictions of the three assays, we define the rank to be $\bar{r}_{ik}$, where $k^{\text{th}}$ is an index over assays. We take the top $10,000$ predicted ranks for each assay and assign $\bar{r}_{ik} = 0$ if the $i^{\text{th}}$ pair is queried but missing from these entries in the $k^{\text{th}}$ assay, and $\bar{r}_{ik} = \min(R_0, \min_{k\neq l}\bar{r}_{il})$ with $R_0 = 2,500$, otherwise. The final rankings are then $\bar{r}' = \sum_k \bar{r}_{ik}/3$. The treatment of the missing entries prioritizes pairs that appear in multiple assays as these pairs are likely more robust than the top predictions in each assay without allowing the absence of a pair in a given assay to increase its average rank. The code for RNM is available at: `https://github.com/spxuw/PPI-Prediction-Project`.

**Other Similarity-Based Methods in the Literature**. There are some additional traditional similarity-based methods in the literature, e.g., the Local Leicht-Holme-Newman Index [27], the Individual Attraction Index [28], the Mutual Information [29], the CAR-Based Indices [30], and the Global Leicht-Holme-Newman Index [27], which can also be used to tackle the link prediction problem (see surveys [31–33] for details).

Most of the similarity and diffusion based link prediction methods are available in the MATLAB implementation of SEAL [34]:
`github.com/muhanzhang/SEAL/tree/master/MATLAB`.

##### B. Probabilistic methods

These methods are based on the concept of maximum likelihood, assuming that real networks have some structure, i.e., hierarchical or community structure. The goal of these algorithms is to select model parameters that can maximize the likelihood of the observed structure.

**Stochastic Block Model**. As one of the most general network models, the stochastic block model (SBM) assumes that nodes are partitioned into groups, and the probability that two nodes are connected depends solely on the groups to which belong [35, 36]. The SBM assumes that a link with higher reliability has higher existent probability, and the reliability of a link is defined as [37]:

$$R_{ij}^{L} = \frac{1}{Z} \sum_{P \in \mathcal{P}} \left( \frac{l_{\sigma_i \sigma_j}^{O} + 1}{r_{\sigma_i \sigma_j} + 2} \right) \exp[-\mathcal{H}(P)], \tag{22}$$

where $\mathcal{P}$ represents the partition space of all possible partitions, $\sigma_i$ is the group to which that node $i$ belongs in the partition $P$, $l_{\sigma_i \sigma_j}^{O}$ is the number of links between groups $\sigma_i$ and $\sigma_j$ in the observed network, $r_{\sigma_i \sigma_j}$ is the maximum possible number of links between them, and the function $\mathcal{H}(P) \equiv \sum_{\alpha \leq \beta} [\ln(r_{\alpha\beta} + 1) + \ln \begin{pmatrix} r_{\alpha\beta} \\ l_{\alpha\beta}^{O} \end{pmatrix}]$, and $Z \equiv \sum_{P \in \mathcal{P}} \exp[-\mathcal{H}(P)]$. In practice, we can use the Metropolis algorithm to sample relevant partitions that significantly contribute to the sum over the partition space $\mathcal{P}$. This approach allows us to calculate the link reliability efficiently.

**Hierarchical Structure Model**. Many real networks have hierarchical structure, which can be represented by a dendrogram $D$. One can assign a probability $p_r$ to each internal node $r$ of $D$ with the connecting probability of a pair of leaves given by $p_{r'}$, where $r'$ is the lowest common ancestor of these two leaves. Denote $E_r$ as the number of edges in the network whose endpoints have $r$ as their lowest common ancestor in the dendrogram $D$. Let $L_r$ and $R_r$ be the number of leaves in the left and right subtrees rooted at $r$, respectively. The likelihood of $D$ associated with a set of probabilities $\{p_r\}$ is, then, given by [38]:

$$\mathcal{L}(D, \{p_r\}) = \prod_{r \in D} p_r^{E_r} (1 - p_r)^{L_r R_r - E_r}. \tag{23}$$

For a specific $D$, the probabilities $\{\overline{p}_r\}$ that maximize $\mathcal{L}(D, \{p_r\})$ are edges between the two subtrees of $r$ that are present in the network:

$$\{\overline{p}_r\} = \frac{E_r}{L_r R_r}. \tag{24}$$

Evaluating the likelihood $\mathcal{L}(D, \{p_r\})$ at this maximum yields

$$\mathcal{L}(D) = \prod_{r \in D} \left[ \bar{p}_r^{\bar{p}_r} (1 - \bar{p}_r)^{1-\bar{p}_r} \right]^{L_r R_r}. \tag{25}$$

One can use the Markov chain Monte Carlo (MCMC) method to sample a large number of dendrograms $D$ with probability proportional to their likelihood $\mathcal{L}(D)$. For each pair of unconnected leaves $i$ and $j$, we calculate the connection probability $p_{ij}$ for each $D$, and then calculate the average $\langle p_{ij} \rangle$ over all the sampled dendrograms. The $\langle p_{ij} \rangle$ value yields the existence probability of the link between nodes $i$ and $j$. For each non-existent link or node pair $(i, j)$, we calculate the average connecting probability $\langle p_{ij} \rangle$ over all sampled dendrograms, with node pairs with highest $\langle p_{ij} \rangle$ being missing links.

**RepGSP**. This method is based on the Graph Signal Processing (GSP) technique [39]. The method consists of a Multiscale Markov Random Field (MRF) of the interactome. MRF is a Markov technique modeling the network's edges based on features associated with the nodes [40]. In order to capture latent information of the network, RepGSP uses signal of graphs (SoGs). The SoGs are vector representations that capture the topological patterns of the network through a function denoted the "Spectral Graph Wavelet Transform". The MRF technique also takes into account the pathways of length 3 between two nodes (i.e., proteins). The method is completely unsupervised and is based only on the structural information of the network.

**Other Probabilistic Models in the Literature**. There are also many probabilistic model based link prediction methods (which typically have high time complexity [41]), such as the Probabilistic Relational Model [42], the Probabilistic Entity Relationship Model [43], and the Stochastic Relational Model [44], as well as many other methods.

##### C. Factorization-based methods

These methods posit that the observed network structure has can be generated from a lower-dimensional *latent space* and use matrix factorization or completion techniques to find a mapping to embed the original network, into a lower dimension such that the similar nodes in the original network tend to have similar latent representation features.

**Matrix Factorization**. Matrix factorization aims to decompose the observed adjacency matrix or node attribute data into two or more matrices through supervised and unsupervised approaches. Denote a data matrix $X$, with $p$ rows and $n$ columns. Each column represents a sample and each row represents a particular feature. The data matrix can be factorized as [32, 45]:

$$X \approx FG^T, \tag{26}$$

where $X \in \mathbb{R}^{p\times n}$, $F \in \mathbb{R}^{p\times k}$, and $G \in \mathbb{R}^{n\times k}$. Matrix $F$ represents the bases of the latent space, and matrix $G$ contains combinations of coefficients of the bases and $k$ is a parameter that specifies the dimension of latent space ($k < n$). The matrix factorization methods can be categorized by the sign constraints on the above three matrices, which are denoted by a subscript. For example, singular value decomposition (SVD): $X_\pm \approx F_\pm G_\pm^T$; Non-negative matrix factorization (NMF): $X_+ \approx F_+ G_+^T$; Semi-NMF: $X_\pm \approx F_\pm G_+^T$. Convex-NMF: $X_\pm \approx F_\pm W_+ G_\pm^T$'. Solving the factorization problem can be modeled as a constrained and potentially regularized least squares optimization problem.

**Low-Rank Matrix Completion**. The goal of matrix completion is to recover a low-rank matrix $L$ from a large matrix $A$, which can be used to infer the missing links of a network. The matrix $L$ can be calculated by solving the convex optimization problem [46]:

$$\min_{L,S} \|L\|_* + \lambda|S|_1, \text{s.t. } A = L + S. \tag{27}$$

Here, $S$ is a sparse matrix. $\|\cdot\|_*$ denotes the nuclear norm (sum of singular values) of a matrix, and $|\cdot|_1$ represents the sum of the absolute values of each matrix element and $\lambda$ is a positive parameter. The matrix $S$ is defined as follows: if two nodes $(i, j)$ are connected in the observed network, then the score $S_{ij} = A_{ij}$; otherwise, $S_{ij} = L_{ij}$.

**GLEE**. The Geometric Laplacian Eigenmap Embedding (GLEE) method uses the Laplacian matrix to find the embedding using (simplex) geometric properties, rather than spectral properties, through leveraging the simplex geometry of the network [47]. The Geometric Laplacian Eigenmap Embedding is available at: `github.com/leotrs/glee`. We used the default parameters in GLEE and dimensionality is set as $d = 128$.

##### D. Diffusion-based methods

For a given network, a diffusion process captures the latent structure in the spread of information over the connections. For example, random walks propagate information by sampling network paths following a stochastic process. Diffusion techniques are used in a variety of settings and are integrated with similarity measures or deep learning methods like the Skip-Gram model [48]. Here, we briefly introduce some widely used diffusion-based algorithms.

**Average Commute Time**. The average commute time index is motivated by a random walk process on the network [7]:

$$s_{ij} = \frac{1}{l_{ii}^{+} + l_{jj}^{+} - 2l_{ij}^{+}}, \tag{28}$$

where $l_{ij}^{+}$ is the entry of the pseudoinverse of the Laplacian matrix $L \equiv D - A$, where $D = \text{diag}\{k_1, k_2, \cdots, k_N\}$ is the degree matrix and $A$ is the adjacency matrix.

**RWR**. This method involves a similarity measure based on a random walk with restart sampling. Random walks with restart are random walks with a restart probability [49].

**SimRank** This method involves a network propagation of node structural context similarity based on object-to-object relationships [50]. SinRank measures similarity of the structural context in which objects occur, based on their relationships with other objects.

**Deepwalk**. DeepWalk uses local information obtained from truncated random walks to learn latent representations by treating walks as the equivalent of sentences, consisting of two main components: random walk sampling and embedidng learning [51]. Once the random walk sequence are sampled, these are fed to a Skip-Gram neural network to learn node embeddings [52].

**LINE**. LINE is a novel network embedding method, with a carefully designed objective function that preserves both the first-order and second-order proximities, which is scale to large directed, undirected and weighted networks [53].

**Node2vec**. node2vec learns a mapping of nodes to a low-dimensional space of features that maximizes the likelihood of preserving network neighborhoods of nodes. By choosing an appropriate notion of a neighborhood, node2vec can learn representations that organize nodes based on their network roles and/or communities to which they belong [54].

**RW2**. This method is divided into two steps: representation learning and classification. In the representation learning step, RW2 is applied to the interactome to generate node embeddings. Since RW2 needs categorical features associated with nodes, each node is enriched by the following labels, node ID and biological properties expressed in terms of GO-Terms (biological functions, biological processes and cellular locations). In the classification step, an XGboost method is trained with the embeddings generated by RW2. The method is supervised and is based on structural and biological information in the network (Level-2).

**DNN+node2vec**. This method consists of two steps. First, it employs node2vec to learn node embeddings [54]. Next, the node embeddings are fed to a deep neural network for PPI prediction. The deep neural network is invariant to the node order in input. The code of DNN+node2vec is available at: `https://github.com/spxuw/PPI-Prediction-Project`.

#### E. Machine Learning method

Machine learning (ML) is a growing field based on learning techniques to extract regular patterns in the given data. Deep learning (DL) is a sub-field of Machine Learning composed of multi-layered neural network methods. Here, we introduce classic machine learning and deep neural network architectures.

**Coding Proteins and K-Nearest Neighbors (Code4+KNN)**. This method first represents each protein sequence as a vector by using local protein sequence descriptors, and then concatenates the information of the two proteins in the pair into a new pairwise feature vector using four "coding" functions. Finally, a KNN method is trained using the pairwise feature vectors as input [55].

**Auto Covariance and Support Vector Machine (AC+SVM)**. This method is a sequence-based method that combines the auto covariance (AC) measure with the support vector machine (SVM) classifier. AC accounts for the interactions between residues of a certain separation distance in the sequence in an attempt to account for local interactions between amino acids. [56].

**Local Protein Sequence Descriptors and Support Vector Machine**. This approach combines local protein sequence descriptors with the SVM classifier. Local descriptors account for the interactions between sequentially distant but spatially close amino acid residues in an attempt to capture multiple overlapping continuous and discontinuous binding patterns within a protein sequence [57].

**Multi-Scale Continuous and Discontinuous feature Representation and Support Vector Machine (MCD+SVM)**. This approach uses a novel Multi-scale Continuous and Discontinuous (MCD) feature representation and Support Vector Machine (SVM). The MCD representation gives adequate consideration to the interactions between sequentially distant but spatially close amino acid residues; thus, it can sufficiently capture multiple overlapping continuous and discontinuous binding patterns within a protein sequence. An effective feature selection method mRMR was employed to construct an optimized and more discriminative feature set by excluding redundant features. Finally, a prediction method is trained and tested using the SVM classifier to predict the interaction probability of protein pairs [58].

**Deep-Forest (GcForest)**. This is a deep-forest-based method for PPIs prediction. First, pseudo amino acid composition (PAAC: an encoding of the amino acids based on their functional role), autocorrelation descriptor (Auto: a measure of how frequently specific amino acids recur), multivariate mutual information (MMI: a method for organizing high-dimensional mutual information terms by the number of variables considered so that the mutual information can be approximated), composition-transition-distribution (CTD. Composition: frequency of amino acids; Transition: frequency of amino acid pairs; distribution: frequency of amino acids in a subdomain in a protein), amino acid composition PSSM (AAC-PSSM: an amino acid position specific substitution matrix constructed from phylogenetic data that measures how frequently the amino acids are substituted in functionally similar genes), and dipeptide composition PSSM (DPC-PSSM: is similar to PSSM, but it is constructed specifically for pairs of amino acids.) are adopted to extract and construct the pattern of PPIs. Secondly, elastic net is used to optimize the initial feature vectors and boost the predictive performance. Finally, the GcForest-PPI method based on deep forest is constructed [59].

**FCTP-WSRC**. Initially, combinations of the F-vector (a fixed length feature vector containing key information), composition (C) and transition (T) are used to map each protein sequence into numeric feature vectors. Principal component analysis (PCA) is employed to reconstruct the most discriminative feature subspaces, which are subsequently used as input in a weighted sparse representation based classification (WSRC) for prediction [60].

**PCA-EELM**. This method involves a novel hierarchical principal component analysis-ensemble extreme learning machine (PCA-EELM) model to predict protein-protein interactions using the information contained in protein primary amino acids sequences. In the proposed method, 11,188 PPIs retrieved from the DIP database were encoded into feature vectors by using four kinds of protein sequences descriptors. Next, the PCA method was employed to construct the most discriminative new feature set. Finally, multiple extreme learning machines were trained and then aggregated into a consensus classifier by majority voting [61].

**DeepPPI**. This method, called DeepPPI (Deep neural networks for Protein-Protein Interactions prediction), employs deep neural networks to learn effectively the representations of proteins from common protein descriptors [62].

**iPPI**. First, iPPI proposes the amino acid properties of PPI closely related protein sequences and the influence of an imbalanced training set on prediction accuracy. Next, it encodes the primary sequence with the properties of 20 amino acids and employs a random undersampling strategy to solve the imbalance issue. In the training phase, it builds a hybrid deep neural network to model the encoded sequence profiles of the bootstrapping-based training datasets [63].

**DPPI**. This method applies a deep, twin-like convolutional neural network combined with random projection and data augmentation to predict PPIs, leveraging existing high-quality experimental PPI data and evolutionary information of a protein pair under prediction [64].

**Deep generative model (DGM)**. The key princple of DGM is to represent the adjacency matrix of a network as an image, then learn hierarchical feature representations of the image by training a deep generative model. Those features correspond to structural patterns in the network at different scales, from small subgraphs to mesoscopic communities. This method does not rely on any domain-specific heuristic and works for general undirected or directed complex networks [65].

**cGAN**. This method consists of training a conditional Generative Adversarial Network (cGAN) method that uses Wasserstein distance-based loss with gradient penalty [66–68]. The method is divided into two steps: representation learning and adversarial learning. In the representation learning step, the authors generate node embeddings by applying node2Vec [54] to the interactome. Next, in the adversarial learning step, they train the cGAN by feeding the node embeddings and adjacency matrices of the subgraphs (generated starting from each node of the interactome) to the cGAN's generator. In the meanwhile, they feed the same matrices given to the generator and the confidence matrices produced by the generator to the cGAN's discriminator that serves as a critic, scoring the performance of the generator. The cGAN architecture does not perform classification directly for the purpose of link prediction, rather to provide feedback for the training of the cGAN's generator. Thus, the generator (an unsupervised method) can be trained using the loss of the discriminator (a supervised method). cGAN might be considered as a self-supervised method, in which only network topology-based features (Level-1) are used. The code of cGAN used to generate the presented results is available at: `https://github.com/spxuw/PPI-Prediction-Project` , and a continuously updated version is available at: `https://github.com/semmelweis-pharmacology/ppi_pred`.

**GraphSAGE**. GraphSAGE is a popular graph representation learning method that uses node feature information to generate node embeddings for previously unseen data. Unlike most of embedding methods that require all nodes in the graph to be present, GraphSAGE trains individual embeddings for each node through learning a function that generates embeddings by sampling and aggregating features from a node's local neighborhood [69].

**SEAL**. SEAL is a representative GCN-based link prediction method, which combines graph structural features, latent features, and explicit features into a single GCN [34]. In particular, the input to GCN is a local subgraph around each target link. Those local subgraphs capture graph structure feature related to link existence. The latent and explicit features can be naturally combined in GCN by concatenating node embedding and node attributes in the node information matrix for each subgraph. We used the the code found at: `github.com/muhanzhang/SEAL/tree/master/Python`. The parameters used for SEAL are number of hop $h = 1$, the maximum node per hop is 50, the maximum training links is 10,000 for HuRI to reduce the memory requirement, while the default parameters are used for the other 4 organism-specific interatomes. The embedding features of each node were obtained by node2vec: `https://github.com/eliorc/node2vec` with default parameters. The dimensionality of HuRI is $d = 4$ and $d = 128$ for other 4 interactomes. The node attributes of each interactome are obtained from the approach proposed in Ref. [60]; this study developed a method, called FCTP-WSRC, to predict PPIs. Here, we used the first 30 dimensions in PCA analysis as the node attributes of each protein. The sequences were downloaded from the Uniprot [70].

**CGNN**. CGNN uses a variational auto-encoder procedure designed to obtain node embeddings using neural networks [71]. The observed variable $A$, evidence feature $X$ and the latent variable $Z$ are connected by the joint distribution representing the probability of $A$ and $X$ with a given $Z$. The neural network is used to learn the joint probability via MCMC through supervised learning.

**SkipGNN**. SkipGNN a graph neural network approach for the prediction of molecular interactions [72]. SkipGNN predicts molecular interactions by not only aggregating information from direct interactions but also from second-order interactions, which the authors call skip similarity. SkipGNN receives neural messages from two-hop neighbors as well as direct neighbors in the interaction network, and non-linearly transforms the messages to obtain useful information for prediction. To inject skip similarity into a GNN, SkipGNN constructs a modified version of the original network, called the skip graph. The SkipGNN is available at: `https://github.com/kexinhuang12345/SkipGNN`. The embedding features of each node were obtained by node2vec: `https://github.com/eliorc/node2vec` with default parameters. The dimensionality is $d = 128$ for all 5 interactomes.

#### II. SUPPLEMENTARY TABLES

| Network | $N$ | $E$ | $\langle k \rangle$ | $d$ | $l$ | $C$ | $\rho$ | $N_{\text{cc}}$ |
|---|---|---|---|---|---|---|---|---|
| *A. thaliana* | 2.774 | 6.205 | 4.474 | 15 | - | 0.049 | 0.00161 | 120 |
| *C. elegans* | 2.528 | 3.864 | 3.057 | 14 | - | 0.019 | 0.00121 | 147 |
| *S. cerevisiae* | 2.018 | 2.930 | 2.904 | 14 | - | 0.0462 | 0.00144 | 185 |
| *Homo sapiens* | 8.272 | 52.548 | 12.705 | 12 | - | 0.059 | 0.00154 | 72 |
| Synthetic | 8.272 | 52.922 | 12.795 | 15 | - | 0.103 | 0.00155 | 8 |
| *A. thaliana* LCC | 2.532 | 6.037 | 4.768 | 15 | 4.657 | 0.051 | 0.00188 | 1 |
| *C. elegans* LCC | 2.214 | 3.659 | 3.305 | 14 | 5.321 | 0.019 | 0.00149 | 1 |
| *S. cerevisiae* LCC | 1.647 | 2.682 | 3.257 | 14 | 5.612 | 0.057 | 0.00198 | 1 |
| *Homo sapiens* LCC | 8.149 | 52.463 | 12.876 | 12 | 3.844 | 0.060 | 0.00158 | 1 |
| Synthetic LCC | 6.597 | 43.695 | 13.247 | 15 | 5.692 | 0.101 | 0.00201 | 1 |

TABLE S1. Network statistics summary. LCC: largest connected component. $N$: number of nodes. $E$: number of edge. $\langle k \rangle$: average degree. $d$: diameter. $l$: characteristic path length. Note that $l$ cannot be calculated for disconnected network. For these cases we use symbol $-$ as value for $l$. $C$: clustering coefficient. $\rho$: edge density. $N_{\text{cc}}$: number of connected components. We used the software Cytoscape to calculate these statistics [73].

| Method | $\rho_0$ | $l_0$ | $f_{\mathrm{sl}}$ |
|---|---|---|---|
| MPS(B&T) | 0.1458 | 2.491 | 0% |
| MPS(T) | 0.0908 | 2.718 | 0% |
| RNM | 0.0943 | 2.348 | 0% |
| SBM | 0.0994 | 1.964 | 0% |
| cGAN | 0.0238 | 2.595 | 99.4% |
| SEAL | 0.0127 | 3.671 | 0% |
| DNN+node2vec | 0.0128 | 3.394 | 0% |

TABLE S2. Network features of the positive PPIs predicted by the top-seven methods. Prior edge density $\rho_0$ measures the edge density of the interactome subgraph induced by the proteins involved in the positive PPIs (among the top-500 predicted PPIs) predicted by a method. Prior average shortest path length $l_0$ is the average shortest path length between the proteins involved in the positive PPIs (among the top-500 predicted PPIs) predicted by a method. Both $\rho_0$ and $l_0$ were calculated in the original interactome (without considering the new PPIs). $f_{\mathrm{sl}}$ represents the fraction of self-loops among the positive PPIs predicted by a method. Note that all the measures presented in this table are based on the positive PPIs within the top-500 predicted PPIs of each method.

| Method | Positive count | Negative cout | Precision |
|---|---|---|---|
| MPS(B&T) | 376 | 54 | 0.874 |
| MPS(T) | 272 | 86 | 0.759 |
| RNM | 274 | 120 | 0.695 |
| SBM | 225 | 167 | 0.574 |
| cGAN | 175 | 164 | 0.516 |
| SEAL | 19 | 332 | 0.054 |
| DNN+node2vec | 10 | 276 | 0.035 |

TABLE S3. Experimental evaluation of the top-seven human PPI prediction methods. Precision is defined as: Precision=Positive count/(Positive count+Negative count).

#### III. SUPPLEMENTARY FIGURES

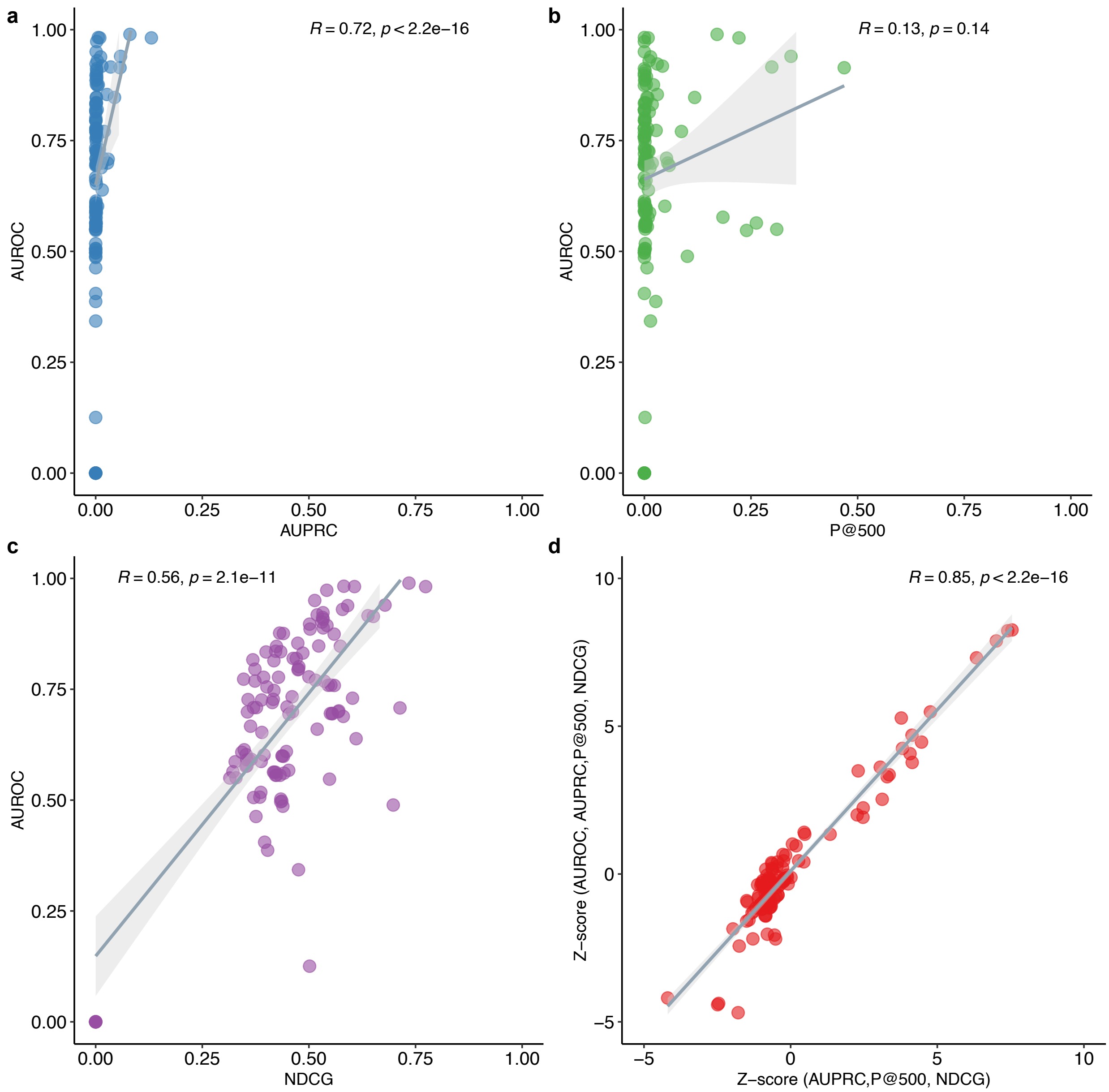


FIG. S1. **Relationship between AUROC and other performance metrics.** Pearson correlation between AUROC and AUPRC (**a**), P@500 (**b**), and NDCG (**c**). **d:** Pearson correlation between Z-scores including AUROC and excluding AUROC. P values were calculated from t-test.

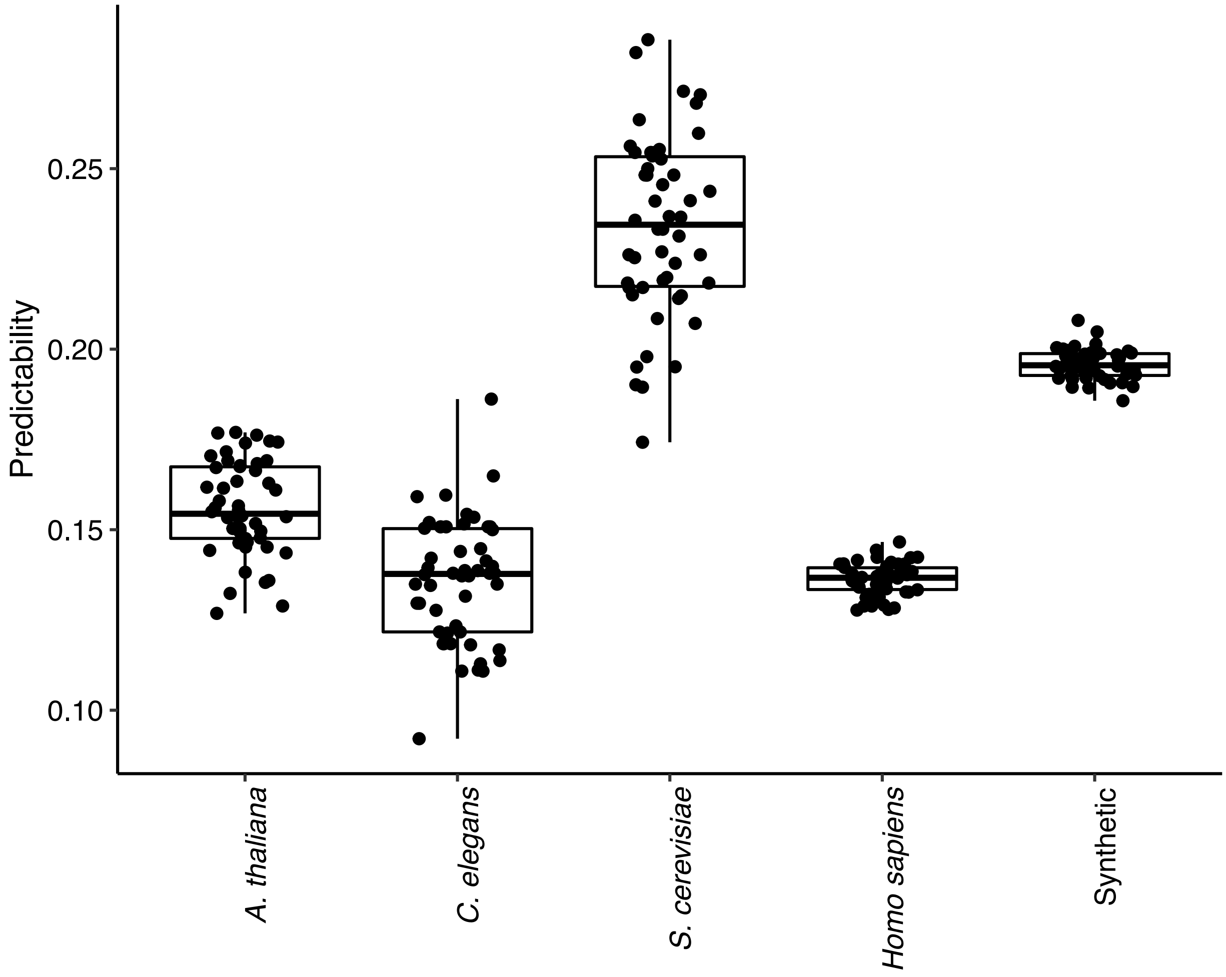


FIG. S2. **Predictability of different interactomes.** Boxplot shows the predictability over 50 different realizations. For each realization, we randomly split the links into the training set (90%), with the remaining 10% as the test set. To quantify the predictability of each interactome, we calculated its structural consistency index $\sigma_c$ based on the first-order perturbation of the interactome's adjacency matrix, using the Matlab implementation of the Structural Perturbation Method (SPM) for link prediction [74]. (Note that here we explicitly considered self-loops in the calculations of $\sigma_{\rm c}$.) See I A for details on SPM.

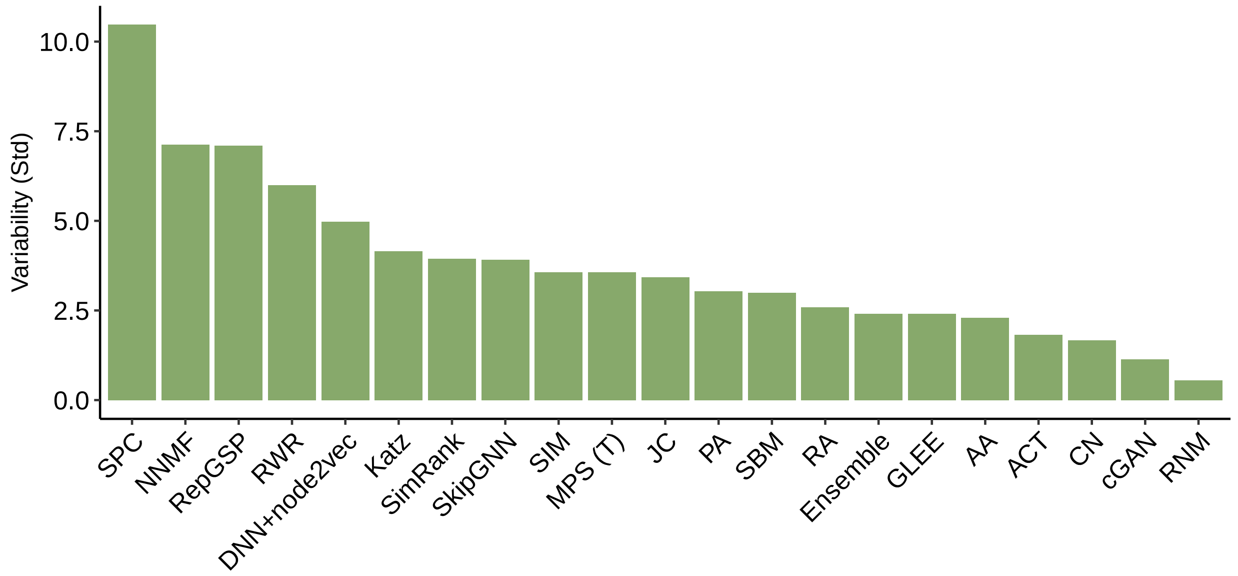


FIG. S3. **Ranking variability of the PPI prediction methods over five interactomes.** The variability of a method is defined as the standard derivation of rankings over five interactomes analyzed in this project. We did not show the variabilities of those methods (in total 3) that were not applied to all the five interactomes.

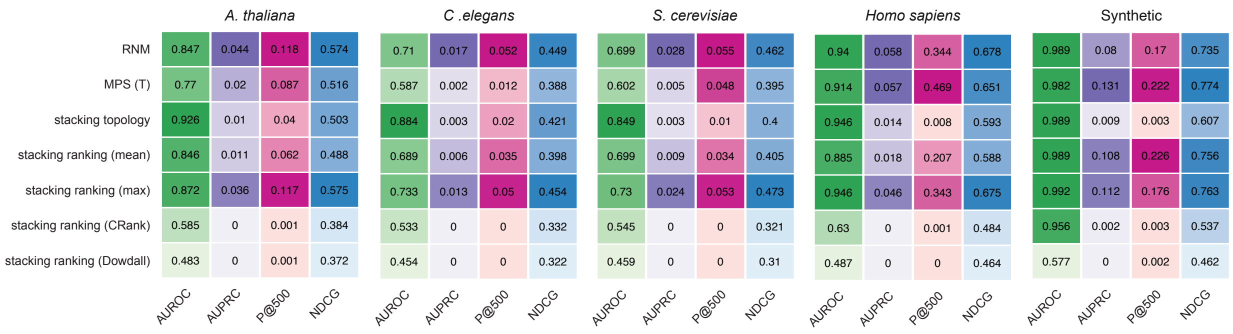


FIG. S4. **Stacking models do not outperform individual methods in any of the five different interactomes.** Stacking topology (Supervised): we stacked 36 individual topological predictors into a single algorithm, then train a supervised classifier to predict the missing links. Stacking ranking (mean) (Unsupervised): we merged the ranking scores of each link from RNM and MPS, into a single value by taking the mean of the two. Stacking ranking (max) (Unsupervised): we merged the ranking scores of each link from RNM and MPS, into a single value by taking the maximum of two. Stacking ranking (CRank): ranking aggregation using CRank algorithm [75]. Stacking ranking (Dowdall): ranking aggregation using Dowdall method [76].

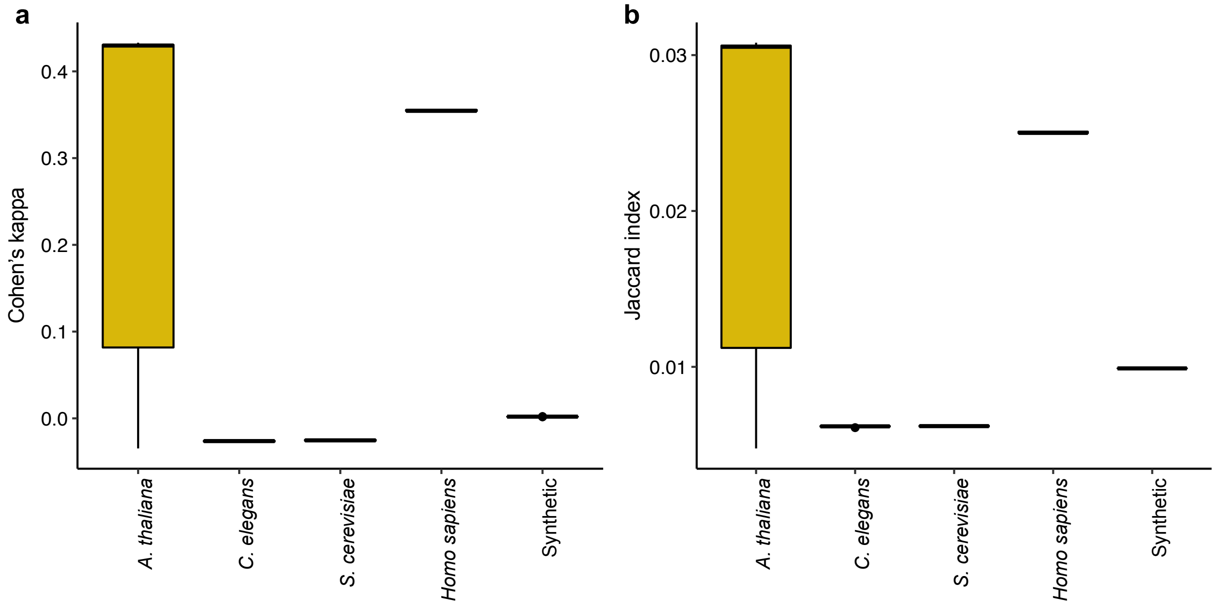


FIG. S5. **Consistency between the ranking from different methods.** **a:** Cohen's kappa coefficient between the ranking scores of all test links under 10-fold cross validation by RNM and MPS(T). **b:** The Jaccard index between the ranking scores of RNM and MPS(T). In the calculation of Cohen's kappa coefficient and the Jaccard index, we set the threshold i.e., the fraction of test links, to be 10%, to dichotomize the ranking scores.

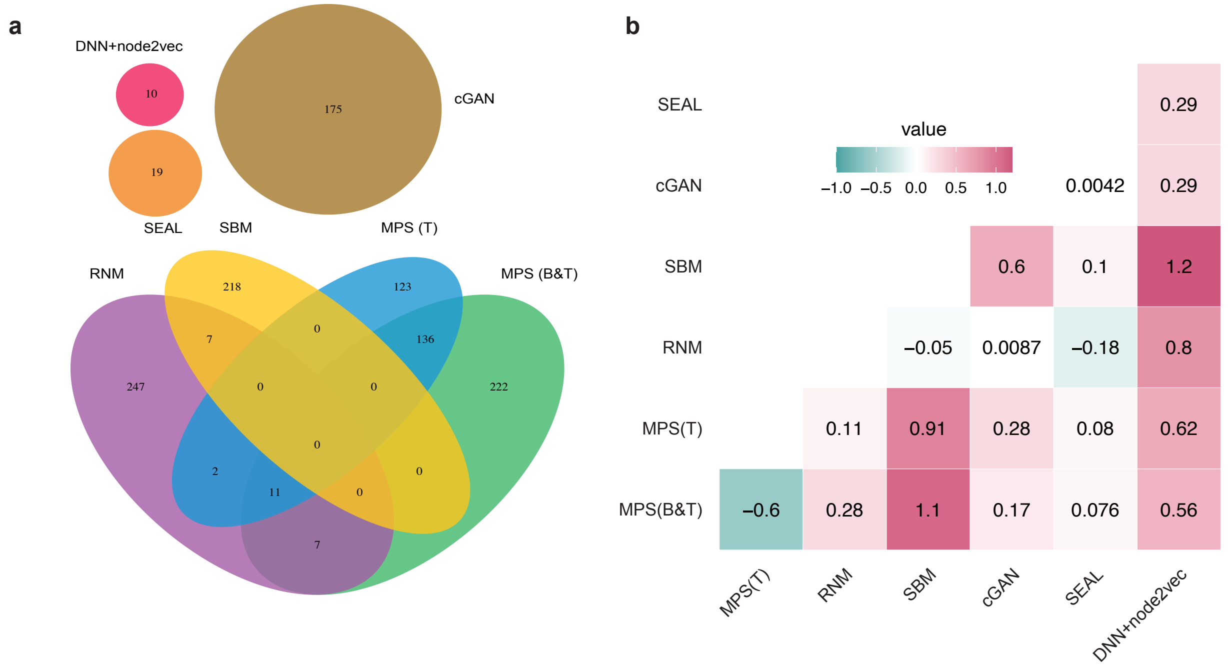


FIG. S6. **Positive PPIs predicted by the top-seven methods do not overlap and represent different areas in the original interactome. a:** Venn diagram for the overlap between the positive PPIs predicted by the top-seven method. **b:** Network separation of proteins involved in the positive PPIs predicted by top-seven methods. The network separation [77] between two sets of proteins is greater than or equal to 0 (red colored cells) if the two protein sets are well separated, if they are overlapping, the separation has a negative value (green colored cells).

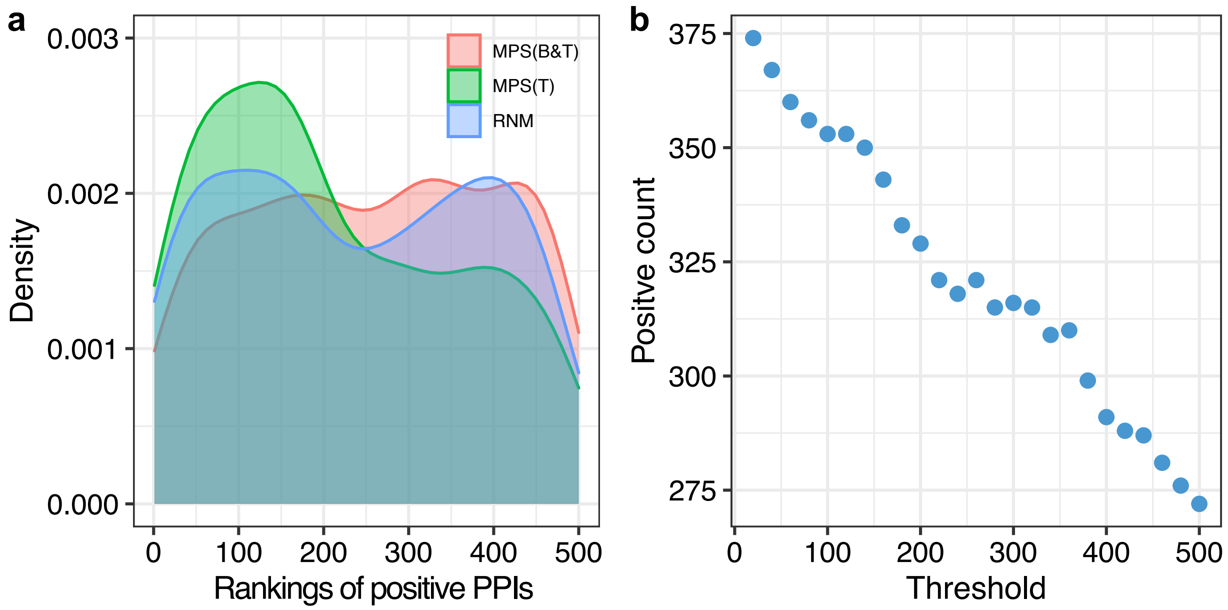


FIG. S7. **Combining the top-500 predicted new PPIs from top-three methods. a:** The ranking distributions (kernel density estimate) of those positive PPIs validated experimentally for each method: MPS(B&T), MPS(T) and RNM. **b**: Number of positive PPIs after combining MPS(B&T), MPS(T) and RNM together: we combined the top-$N_k$ PPIs predicted by MPS(B&T) and top-$[(500 - N_k)/2]$ from MPS(T) and RNM where $0 \leq N_k \leq 500$ is a parameter used to tune the ratio of PPIs predicted by different methods. Note that during the combining process, we ensured that those PPIs predicted by multiple methods appear only once in the top-500 PPI list.

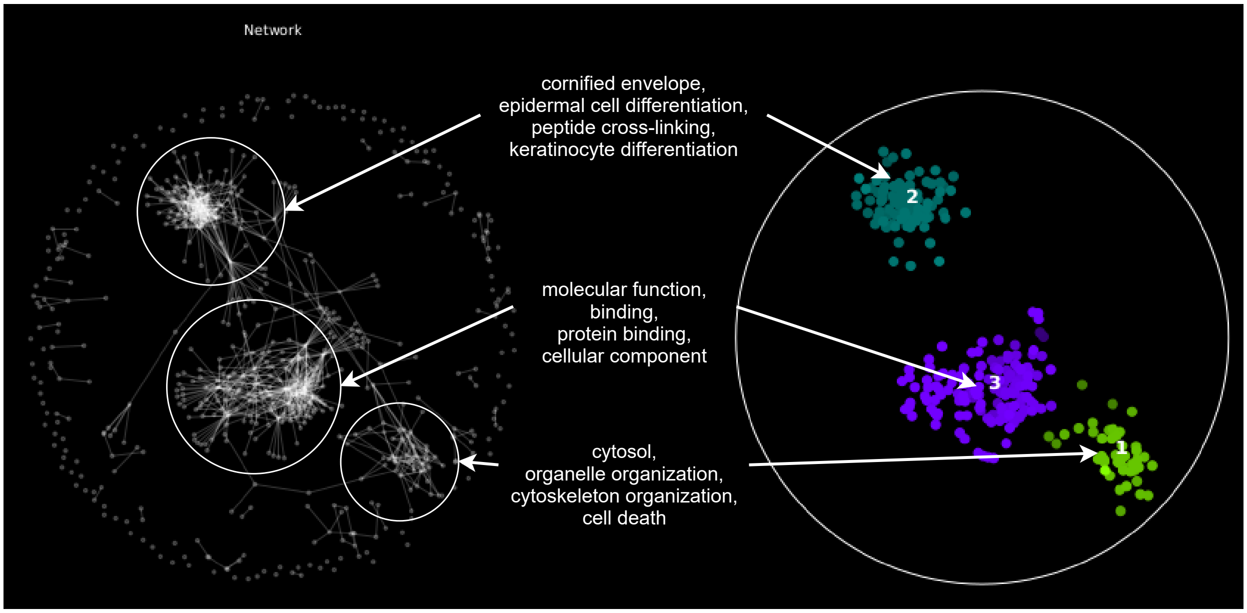


FIG. S8. **Functional relationships among the new human PPIs discovered in this project.** Left: This network consists of all 1,177 new human PPIs predicted by the top-seven methods and validated by Y2H assays. Right: functional modules discovered by SAFE [78]. Gene Ontology [79] (GO) terms for each gene were extracted from FuncAssociate [80]. The neighborhood radius is set to be 0.15 in SAFE. Note that the PPIs in the first and the second modules are mostly predicted by MPS(B&T) and MPS(T) but the rest of the PPIs are predicted by the other methods.

[1] J. De Las Rivas and C. Fontanillo, Protein–protein interactions essentials: key concepts to building and analyzing interactome networks, PLoS Comput Biol **6**, e1000807 (2010).

[2] K. Titeca, I. Lemmens, J. Tavernier, and S. Eyckerman, Discovering cellular protein-protein interactions: Technological strategies and opportunities, Mass Spectrometry Reviews **38**, 79 (2019).

[3] A. R. Mashaghi, A. Ramezanpour, and V. Karimipour, Investigation of a protein complex network, The European Physical Journal B-Condensed Matter and Complex Systems **41**, 113 (2004).

[4] M. Al Hasan, V. Chaoji, S. Salem, and M. Zaki, Link prediction using supervised learning, in *SDM06: Workshop on link analysis, counter-terrorism and security* (2006).

[34] M. Zhang and Y. Chen, Link prediction based on graph neural networks, in *NeurIPS* (2018).

[49] H. Tong, C. Faloutsos, and J.-Y. Pan, Fast random walk with restart and its applications, in:Proceedings of the 6th International Conference on Data Mining (2006).

[50] G. Jeh and J. Widom, Simrank: a measure of structural-context similarity, in *Proceedings of the eighth ACM SIGKDD international conference on Knowledge discovery and data mining* (ACM, 2002) pp. 538–543.

[51] B. Perozzi, R. Al-Rfou, and S. Skiena, Deepwalk: Online learning of social representations, in *Proceedings of the 20th ACM SIGKDD international conference on Knowledge discovery and data mining* (ACM, 2014) pp. 701–710.

[52] T. Mikolov, I. Sutskever, K. Chen, G. S. Corrado, and J. Dean, Distributed representations of words and phrases and their compositionality, in *Advances in neural information processing systems* (2013) pp. 3111–3119.

[53] J. Tang, M. Qu, M. Wang, M. Zhang, J. Yan, and Q. Mei, Line: Large-scale information network embedding, in *Proceedings of the 24th International Conference on World Wide Web*

(International World Wide Web Conferences Steering Committee, 2015) pp. 1067–1077.

[59] B. Yu, C. Chen, Z. Yu, A. Ma, B. Liu, and Q. Ma, Prediction of protein-protein interactions based on elastic net and deep forest, BioRxiv  (2020).

[71] T. Yamamoto, Crystal graph neural networks for data mining in materials science (2019).
